# Symmetry in frontal but not motor and somatosensory cortical projections

**DOI:** 10.1101/2023.06.02.543431

**Authors:** Andrew E. Papale, Madhumita Harish, Ronald F. Paletzki, Nathan J. O’Connor, Brian S. Eastwood, Rebecca P. Seal, Ross S. Williamson, Charles R. Gerfen, Bryan M. Hooks

**Affiliations:** Laboratory of Systems Neuroscience, NIMH, Bethesda, MD; Department of Neurobiology, University of Pittsburgh School of Medicine, Pittsburgh, PA; MBF Bioscience, Williston, VT; Department of Psychiatry, University of Pittsburgh School of Medicine, Pittsburgh, PA; Department of Otolaryngology, University of Pittsburgh School of Medicine, Pittsburgh, PA

**Author notes:** Corresponding authors (BMH); (CRG); (AEP), Correspondence: Dr. Bryan M. Hooks, 200 Lothrop Street BSTWR Suite W1458 Pittsburgh, PA 15213.

## Abstract

Neocortex and striatum are topographically organized for sensory and motor functions. While sensory and motor areas are lateralized for touch and motor control, respectively, frontal areas are involved in decision making, where lateralization of function may be less important. This study contrasted the topographic precision of cell type-specific ipsilateral and contralateral cortical projections while varying the injection site location in transgenic mice of both sexes. While sensory cortical areas had strongly topographic outputs to ipsilateral cortex and striatum, they were weaker and not as topographically precise to contralateral targets. Motor cortex had somewhat stronger projections, but still relatively weak contralateral topography. In contrast, frontal cortical areas had high degrees of topographic similarity for both ipsilateral and contralateral projections to cortex and striatum. Corticothalamic organization is mainly ipsilateral, with weaker, more medial contralateral projections. Corticostriatal computations might integrate input outside closed basal ganglia loops using contralateral projections, enabling the two hemispheres to act as a unit to converge on one result in motor planning and decision making.

**Significance Statement:** Each cerebral hemisphere is responsible for sensation and movement of the opposite side of the body. Many axonal projections cross the midline to target contralateral areas. Crossed corticocortical, corticostriatal, and corticothalamic projections originate from much of neocortex, but how these projections vary across cortical regions and cell types is unknown. We quantify differences in the strength and targeting of ipsilateral and contralateral projections from frontal, motor, and somatosensory areas. The contralateral corticocortical and corticostriatal projections are proposed to play a larger role in frontal areas than in sensory or motor ones as a circuit basis for unifying computation across hemispheres in motor planning, while contralateral connectivity plays a smaller role in sensory and motor processing.

## Introduction

Somatotopic organization of primary motor (M1) and somatosensory (S1) cortices has been established by intracortical stimulation after being predicted by the Jacksonian march of seizures (Penfield & Boldery, 1937; York & Steinberg, 2011). Topography extends to other primary sensory areas, including visual and auditory cortices (Drager, 1975; Merzenich & Brugge, 1973; Morel et al., 1993), and is maintained in higher order sensory areas (Carvell & Simons, 1986; Felleman & Van Essen, 1991; Wang & Burkhalter, 2007). Corticocortical projections, such as from S1 to secondary somatosensory cortex (S2), as well as across modalities, as from S1 to M1, maintain some topography: The S1 to S2 projection is anisotropic, favoring integration across whisker rows (Hoffer et al., 2003), while the S1 to M1 projection is elongated across a broad anterior-posterior swath of M1 (Mao et al., 2011; Rocco-Donovan et al., 2011). Quantification of the precision of somatotopic organization has been a challenge in motor areas, where somatotopy is less precise than in S1 (Hooks et al., 2018). Precision is limited by the microstimulation methods used to quantify somatotopy, which vary with threshold current used (Hooks et al., 2011; Tennant et al., 2011) and current spread (Histed et al., 2012). Anatomical methods lose precision as tracer or virus spreads at the injection site (Bohland et al., 2009; Oh et al., 2014; Zingg et al., 2014). The axes for topographic organization in secondary motor areas are less clear if these lack a body map but instead account for more abstract variables such as which of several motor plans to select. Thus, topographic organization can be less strongly tied to a specific body location. These areas send projections across the midline to the corresponding contralateral cortex, suggesting an important role in coordinating perception and movement.

Corticostriatal projections are also topographically organized, but cortical output from any given cortical area projects somewhat broadly, overlapping with the output from other cortical areas and not perfectly segregated into loops (Alloway et al., 2000; Hintiryan et al., 2016). These projections originate mainly from layer 5, with some contribution from layer 2/3 (McGeorge & Faull, 1989). Striatal regions integrate input from functionally related cortical areas. The main features are the convergence of excitatory cortical (and some thalamic) input from a large number of afferents onto the striatal projection neurons (SPNs) (Lee et al., 2020). Basal ganglia projections are organized to form loops from cortex through basal ganglia and thalamus that are parallel and functionally segregated (Alexander et al., 1986). These loops include inputs from related cortical areas (Middleton & Strick, 2000). These areas may be reciprocally connected, as somatotopically congruent regions of M1 and S1 (Alloway et al., 2006; Hooks et al., 2018) or grasp-related regions across the cortex (Gerbella et al., 2016), but also include non-connected cortical areas (Choi et al., 2017; Selemon & Goldman-Rakic, 1985). These loops are not exclusively concerned with motor control, but perform similar computations on incoming cortical information from frontal, premotor, motor, and temporal inputs (Middleton & Strick, 2000). Thus, these loops are “globally parallel” but “locally overlapping” (Haber, 2003; Jarbo & Verstynen, 2015). Striatal input in non-human primate is dominated by ipsilateral input. There is not strong evidence for left/right asymmetry in ipsilateral corticostriatal projections (McGuire et al., 1991a, 1991b). Some cortical areas send contralateral projections, including parietal, motor, and frontal areas (Alloway et al., 2006; Borra et al., 2022; Jones et al., 1977; Kunzle, 1975; McGuire et al., 1991b). As with ipsilateral projections, there is overlap of striatal targeting from related cortical areas.

Contralateral corticothalamic projections have received less attention, in part because they are weaker than ipsilateral ones, though this may depend on cortical location (Alloway et al., 2008; Hooks et al., 2013; Szczupak et al., 2021). Further, the importance in humans is unclear, as absence of an intrathalamic adhesion through which the fibers would cross does not result in an apparent deficit (Olry & Haines, 2005).

Here, we study the ipsi- and contralateral corticocortical, corticostriatal, and corticothalamic projections of three major cell types in frontal, primary motor, and primary somatosensory cortex using transgenic mice to label intratelencephalic (IT-type), pyramidal tract type (PT-type), and corticothalamic (CT-type) neurons (Gerfen et al., 2013). We find significant symmetry in frontal corticocortical and corticostriatal connections, which may play a role in motor planning and decision making.

## Methods

### Injections

Surgical and experimental procedures conformed to National Institutes of Health guidelines for mice and were approved by the Institutional Animal Care and Use Committees of University of Pittsburgh and Janelia Research Campus. Tlx3_PL56 mice (N=33), Sim1_KJ18 mice (N=22), and Ntsr1_GN220 mice (N=6) from the GENSAT BAC Cre-recombinase driver line were used to trace corticocortical and corticostriatal projections. Mice of both sexes were injected at postnatal day ∼P35 and sacrificed after 2-3 weeks of expression. Stereotaxic injections targeted the left hemisphere, with injection sites covering frontal, motor, and somatosensory. Injections in other cortical areas were excluded. 30 nL per injection site of AAV-flex-XFPs were injected at each of two cortical depths (∼0.5 and 0.8 mm) using a custom positive displacement injector via a pulled borosilicate glass micropipette (Hooks et al., 2018). The generic AAV-flex-XFP refers to several tracing viruses used (see Table 1), including EGFP, tdTomato, and mRuby2-based spaghetti monster fluorescent proteins (smFPs) (Viswanathan et al., 2015).

**Table 1.** Viral reagents for tracing.

| Construct name | Addgene # | Addgene name |
| --- | --- | --- |
| AAV2/1-CAG-flex-EGFP | 51502 | pCAG-FLEX-EGFP-WPRE |
| AAV2/1-CAG-flex-tdTomato | 51503 | pCAG-FLEX-tdTomato-WPRE |
| AAV2/1-CAG-flex-GFPsmFP-FLAG | 59756 | pCAG-smFP-FLAG |
| AAV2/1-CAG-flex-GFPsmFP-Myc | 59757 | pCAG-smFP-Myc |
| AAV2/1-CAG-flex-GFPsmFP-V5 | 59758 | pCAG-smFP-V5 |
| AAV2/1-CAG-flex-GFPsmFP-HA | 59759 | pCAG-smFP-HA |
| AAV2/1-CAG-flex-mRuby2smFP-FLAG | 59760 | pCAG-mRuby2-smFP-FLAG |
| AAV2/1-CAG-flex-mRuby2smFP-OLLAS | 59761 | pCAG-mRuby2-smFP-OLLAS |

### Histology, staining, and imaging

Mice were transcardially perfused with phosphate-buffered saline (PBS) followed by 4% paraformaldehyde in PBS and postfixed overnight. Brains were then transferred to 20% sucrose in PBS. Brains were sectioned at 80 µm. EGFP, tdTomato, and smFPs were immunoamplified. Neurotrace Blue (1:100) was used as a structural marker for alignment and registration.

Sections were imaged using Neurolucida (v2017, MBF Bioscience, Williston, VT) on a Zeiss Axioimager (Zeiss, Oberkochen, Germany) equipped with 10x objective, Ludl motorized stage and a Hamamatsu Orca Flash 4.0 camera (Hamamatsu, Hamamatsu City, Japan). Brain sections were tiled images of the average of ∼100-200 tiles. Each tile consisted of image stacks collected in 10 µm steps. A single 3D image was first generated and images were collapsed to a single plane using a DeepFocus algorithm implemented in Neurolucida (Gerfen et al., 2013; Paletzki & Gerfen, 2015).

### Whole brain reconstruction, annotation, and registration

Stacks of brain images were aligned to a standard coordinate system using BrainMaker software (MBF Bioscience, Williston, VT) as previously described (Eastwood et al., 2019; Tappan et al., 2019). Briefly, serially-reconstructed brains contained 10 µm isotropic voxels (782x1086x1242) and were registered to the annotated Allen Mouse Common Coordinate Framework (CCFv3; http://connectivity.brain-map.org) (Wang et al., 2020). All brains were registered to this framework using Neurotrace Blue was used as the structural marker for registration. This required a two-stage registration process. The first stage constructed an average reference space from NeurotraceBlue images (the average of brains sectioned, mounted, and stained with Neurotrace Blue; 78 individual brains). Multiple resolution registration optimized 12 parameters of a 3D affine transform to minimize a normalized correlation metric between each brain and the template image. The reference template was then updated by resampling all individual brains with their respective affine transforms and computing a voxel-wise weighted average. A second pass registered each individual brain to the new template, updating the individual transforms. This process repeated until the template image stabilized.

The second stage registered the average reference image to the Allen CCFv3 (based on 1675 brains visualized with 2--photon auto-fluorescence (Kuan et al., 2015)). 300 unique landmarks were identified and positions of the landmarks were used to construct a nonlinear transform that models deformation of a uniform mesh grid with B-splines. The result is an average reference image and its spatially aligned annotation volume that constitutes the average reference atlas. Individual brains here were registered using NeurotraceBlue with the average reference space by adjusting parameters of a 3D affine and 3D nonlinear B-spline transform to minimize a normalized correlation metric.

For quantification of injection site location, tiled images were imported into Neurolucida software (MBF Bioscience, Williston, VT) and soma locations were annotated using automated object detection with manual supervision. Nearest neighbor interpolation of the average reference space volume at the mapped positions provided the anatomical region assignment for each cell. Coordinates of the CCFv3 for structures of interest (cortex and striatum) were used to identify voxels for quantification. These were divided into left and right hemispheres to distinguish ipsilateral and contralateral sides.

### Data analysis

Aligned brain images were downsampled to 50 µm isotropic voxels (156x217x248) using custom routines in FIJI software. The annotated Allen Mouse CCF was also used at 10 µm and downsampled to 50 µm. The annotation was used to assign voxels to a given brain region (ipsilateral or contralateral striatum, for example). Both 10 µm and 50 µm images were converted from tifs into .mat files in Matlab (Mathworks, Natick, MA) for analysis with custom routines. Soma locations were similarly imported to Matlab. Differences in voxel intensity and the intensity ratio between ipsi- and contralateral sides (Fig. 3D-E and Fig. 7D-E) were tested by one way ANOVA followed up by Tukey-corrected pairwise t-tests. Differences in the slope (versus injection site offset, Fig. 3G-I and Fig. 7G-I) were assessed by two-way ANCOVA to examine the effects of region and ipsi/contra target sites on correlations (PCC), after controlling for distance. Follow up analyses were performed with Bonferroni correction. Statistics were done in R or Matlab.

### Data availability statement

The data and images used in this study are available upon request. Aligned images in 10 and 50 μm voxels for all brains, cell soma locations, and the corresponding masks used to identify brain regions (striatum and cortex) are available on request as well as the custom Matlab code used for data analysis. Original images of whole brains are freely available online at http://gerfenc.biolucida.net/link?l=Jl1tV7.

## Results

### Examining a dense database of crossed corticocortical and corticostriatal projections

Neocortex contains three main subcortical projection neuron types, intratelencephalic (IT-type) and pyramidal tract type (PT-type) in layer 5, and corticothalamic type (CT-type) in layer 6. IT-type neurons project to both ipsi- and contralateral cortex and striatum. PT-type neurons project to ipsilateral striatum only (Hooks et al., 2018), but target thalamus (as well as brainstem and spinal targets) along with CT-type neurons. To generate the image database (Hooks et al., 2018), a Cre-driver mouse line selective for the specific cell type of interest (IT-type neurons, for example) (Gerfen et al., 2013) was injected with AAV expressing Cre- dependent tracers, including GFP, td-tomato, and smFPs (Table 1) (Viswanathan et al., 2015).

Injections were made in a range of locations, including frontal, motor, and somatosensory cortex (Fig. 1). The anterior dorsal surface of mouse cortex has many different names including secondary motor (MOs (Wang et al., 2020; Zingg et al., 2014)) or M2 (Hooks et al., 2013)), anterior lateral motor (ALM (Komiyama et al., 2010)), lateral and medial agranular (AGl and AGm (Li & Waters, 1991)) and FrA (Paxinos & Franklin, 2004). Here, we refer to regions of anterior cortex including M2 and ALM as frontal (Hooks et al., 2018) instead of simply premotor, to differentiate them from primary motor cortex, as these areas play a role in not only motor planning, but also in decision-making (Hamaguchi et al., 2022; Chen et al., 2024; Guo et al., 2014; Li et al., 2016). M2 is the more medial of the two regions, reciprocally connected with whisker M1 (Hooks et al., 2013; Zingg et al., 2014; see coordinates in Hooks et al., 2018). ALM is lateral to M2, and is reciprocally connected to more lateral regions in primary motor and somatosensory cortex (Hooks et al., 2018), though it clearly plays a functional role in planning future volitional movements (Guo et al., 2014). We also noted that these frontal areas target different thalamic nuclei than primary motor areas.

**Figure 1.**
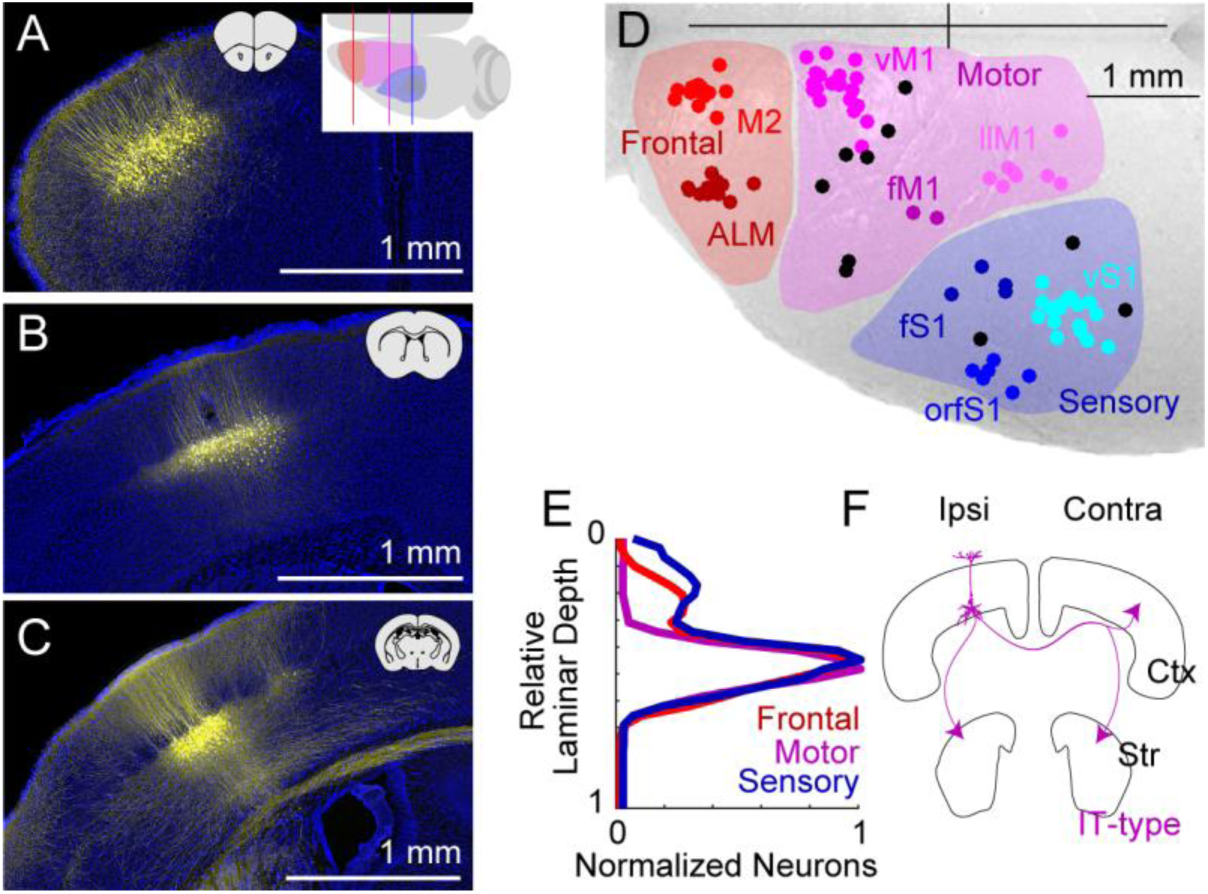
Mapping layer 5 corticostriatal output from sensory and motor cortex. Injection site of AAV-DIO-XFP into frontal (A), motor (B), and sensory cortex (C) of Tlx3_PL56-Cre+ mice. Inset of coronal brain slices shows approximate location. In (A), top-down cartoon shows approximate planes of section. (D) Center of mass of all injection sites in sensory (blue, N=27), motor (purple, N=30), and frontal (red, N=25) cortex. Subdivisions within areas (M2, ALM, vM1, fM1, llM1, vS1, fS1, and orfS1) are indicated in different colors, with uncategorized injection sites in black. Bregma is indicated as the cross at top right. (E) Quantification of relative depth of labeled Cre+ cortical neurons in frontal (red), motor (purple), and sensory (blue) cortex. (F) Cartoon of output targets of IT-type neurons.

Each brain was reconstructed from tiled fluorescence images (Paletzki & Gerfen, 2015). These were registered to the Allen Common Coordinate Framework (CCFv3) (Wang et al., 2020) using BrainMaker software (MBF Bioscience). Alignment precision was about ∼50-70 µm (Hooks et al., 2018). Placing all voxels and detected objects, such as somata, in a common coordinate space enabled quantitative comparisons of brain regions across different animals.

The original images are freely available online at: http://gensatcrebrains.biolucida.net/images/?page=images&selectionType=collection&selectionId=32 (web-based) or can be viewed using the free Biolucida viewer software downloaded from: http://gerfenc.biolucida.net/link?l=Jl1tV7.

Each injection was quantified in two ways. The first relied on detected cell bodies at the injection site. Injection site location was quantified as the center of mass of all infected somata. The somata of L5 IT-type neurons were detected and marked in Neurolucida. Their relative laminar depth was largely restricted to L5 (Fig. 1A-C), though a small number of L2/3 neurons could be seen, which was greatest in somatosensory injections (Fig, 1E). The injection site center of mass was calculated in coordinates registered to the Allen CCFv3, and plotted from a dorsal viewpoint (Fig.1D). The location was used to cluster injection sites into frontal (red), motor (purple), and somatosensory (blue) (Hooks et al., 2018). This clustering corresponded well to existing division of the neocortex (Oh et al., 2014; Zingg et al., 2014) into primary somatosensory (SSp), primary motor (MOp), and secondary motor (MOs) regions. Furthermore, motor and frontal areas correspond to low threshold regions for evoking somatotopically organized body movements in rodent neocortex (Brecht et al., 2004; Hooks et al., 2011; Li & Waters, 1991; Tennant et al., 2011). Injection sites could be further subdivided within general areas into subdivisions, such as whisker, forelimb, and orofacial regions of primary somatosensory cortex (vS1, fS1, and orfS1) which correspond to the CCFv3 regions SSp-bfd, SSp-ul, and SSp-n. Motor subdivisions (whisker, forelimb, and lower limb: vM1, fM1, llM1) were also made, consistent with microstimulation maps and previous analysis of the injection sites (Hooks et al., 2018). Frontal injections targeted the more medial M2 (Hooks et al., 2013) and the more lateral ALM (Komiyama et al., 2010). These did not correspond to subregions within the CCFv3 because the primary and secondary motor areas MOp and MOs are not somatopically divided in that atlas. Some sites (black dots in Fig. 1D, 9D, and 11D) were not categorized with these subdivisions and were only included when the analysis covered the general somatosensory, motor, or frontal area (such as Fig. 3).

The second quantification of each injection compared projections from these injection sites. Thus, this relies on detecting suprathreshold axonal fluorescence from injections. A threshold was applied to aligned images to eliminate 99% of background voxels in target regions where no IT-type projections were expected (thalamus and hippocampus) for the fluorescence channel associated with a given injection. Aligned images are presented throughout using 50 μm-size voxels (50 μm x 50 μm x 50 μm).

### High correlation of projection targeting in IT-type corticocortical projections

IT-type neurons project to four major regions: ipsilateral and contralateral cortex and ipsilateral and contralateral striatum (Fig. 1F). Some individual IT-type neurons collateralize to all four areas, while others show different patterns (Economo et al., 2016; Winnubst et al., 2019). These projections to both cortex and striatum suggest two straightforward comparisons across the midline. Examination of contralateral corticocortical projections suggests differences in the symmetry of these projections across the surface of cortex (Fig. 2). Frontal cortex injections had relatively strong axonal fluorescence in contralateral cortex (Fig. 2A-D), while this fluorescence became weaker in motor injections (Fig. 2E-H) and weaker still in somatosensory ones (Fig. 2I-L). This is not because somatosensory neurons are incapable of making long-range projections, as they clearly project well to topographically appropriate targets in ipsilateral motor cortex (Fig. 2M-P). Further, they make strong projections to ipsi- and contralateral entorhinal cortex (Fig. 2Q-X).

**Figure 2.**
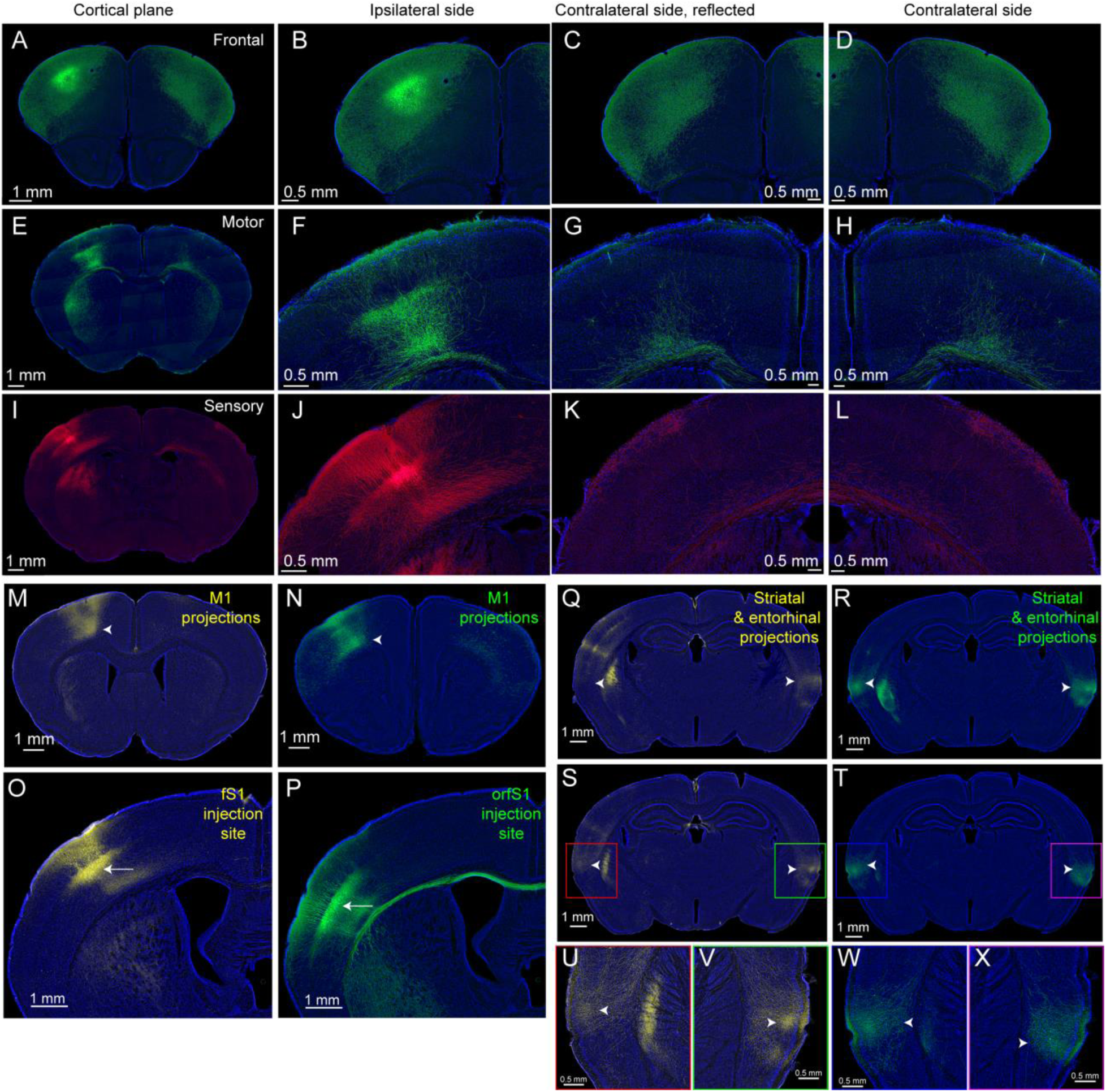
Ipsi- and contralateral projections of IT-type neurons in frontal, motor, and somatosensory cortex. (A-D) Coronal images showing injection site of AAV-DIO-XFP into frontal cortex (ALM), with ipsi- (B) and contralateral (D) cortex presented for comparison. Reflected image of contralateral side (C) shown for comparison. (E-H) Similar presentation for motor cortex (fM1) injection. (I-L) Similar presentation for somatosensory cortex (vS1) injection. (M-X) Comparisons of projections from different regions within S1. (M-N) M1 projections from fS1 (yellow) and orfS1 (green). (O-P) Injection sites. (Q-X) Striatal and entorhinal projections, with higher resolution images (U-X). Arrows show cell bodies; arrowheads show axonal projections.

How accurately do these representative images convey the information contained in the whole dataset? To make quantitative comparisons, analysis proceeded as follows. Both the injection site location and the suprathreshold voxel intensity were used. For the injection sites, the distance between the center of mass of each injection was calculated as a distance offset (in millimeters, Fig. 3A). Because brains are aligned to standard coordinates, injection site offset is used as an estimate of the expected difference in topographic location between two given injections. For axonal projections, all suprathreshold voxels for each injection are used. Pairs of injections are compared on a voxel-by-voxel basis using voxels that are suprathreshold for both injections. Voxels in ipsilateral and contralateral areas are compared based on regional identity in the Allen CCFv3, where each voxel is assigned to a specific brain region or cortical area.

These areas include MOp and MOs for primary and secondary motor cortex as well as SSp-ul, SSp-bfd for the primary somatosensory cortex for upper limb and barrel field respectively. Depending on the analysis, cortical areas are pooled or analyzed by individual topographic areas. The Pearson Correlation Coefficient (PCC) is then computed based on voxel intensities (in arbitrary units, AU) in defined areas and used to assess the relationship for each pair of injections (Fig. 3B).

Example images of the ipsilateral cortex are shown for two aligned brains with somatosensory injections from anterior to posterior, spaced about 0.75 mm. Images range from frontal cortex (∼1.75 mm anterior to bregma to ∼1.25 mm posterior; Fig. 3C). Note the S1 axons in M1 (upper panels) and the brighter injection sites (lower panels). The injection site is always in left cortex, though differences in dominant paw (not assessed here) might affect lateralization of function (Deemyad et al., 2022). Injections in S1 were generally stronger with brighter total voxel intensity when compared to M1 and frontal injections (Fig. 3D). However, the contralateral projections were relatively brighter in frontal injections based on the ratio of contralateral/ipsilateral intensity (Fig. 3E).

Injection sites in nearby locations in aligned brains showed high degrees of correlation in the targeting of their ipsilateral corticocortical projections (Fig. 3G). Examining the ipsilateral somatosensory plot (royal blue), injections with nearly no offset have greater than 0.6 correlation (linear fit near 0 offset). This drops quickly with even small offsets. Motor correlations (purple) follow a similar pattern, though the peak correlation is lower and the slope is not as steep. Frontal cortex (red) also shows a similar pattern to somatosensory, with very high (>0.7) nearby correlation, and very steep decline. Thus, ipsilateral corticocortical projections reliably target similar cortical areas across different animals. What about contralateral cortical projections? These also showed some correlation that varied with injection site offset (Fig. 3H). But the pattern differed from ipsilateral cortex. Specifically, somatosensory cortex had correlations (light blue) that were lower than ipsilateral ones (Fig. 3I). Similarly, contralateral motor cortex projections (light purple) were not as well somatotopically organized as ipsilateral ones, though in general not as poor as those in somatosensory areas. Frontal cortex injections (pink) had quite strong corticocortical correlations for contralateral projections. The correlation data was also assessed in a similar manner for subdivisions of somatosensory, motor, and frontal cortex (vS1, fS1, orfS1, vM1, fM1, llM1, M2, and ALM; Fig. 3F). Notably, vS1 correlations were lower than other topographic areas of sensory cortex and llM1 injections were lower than other motor areas.

**Figure 3.**
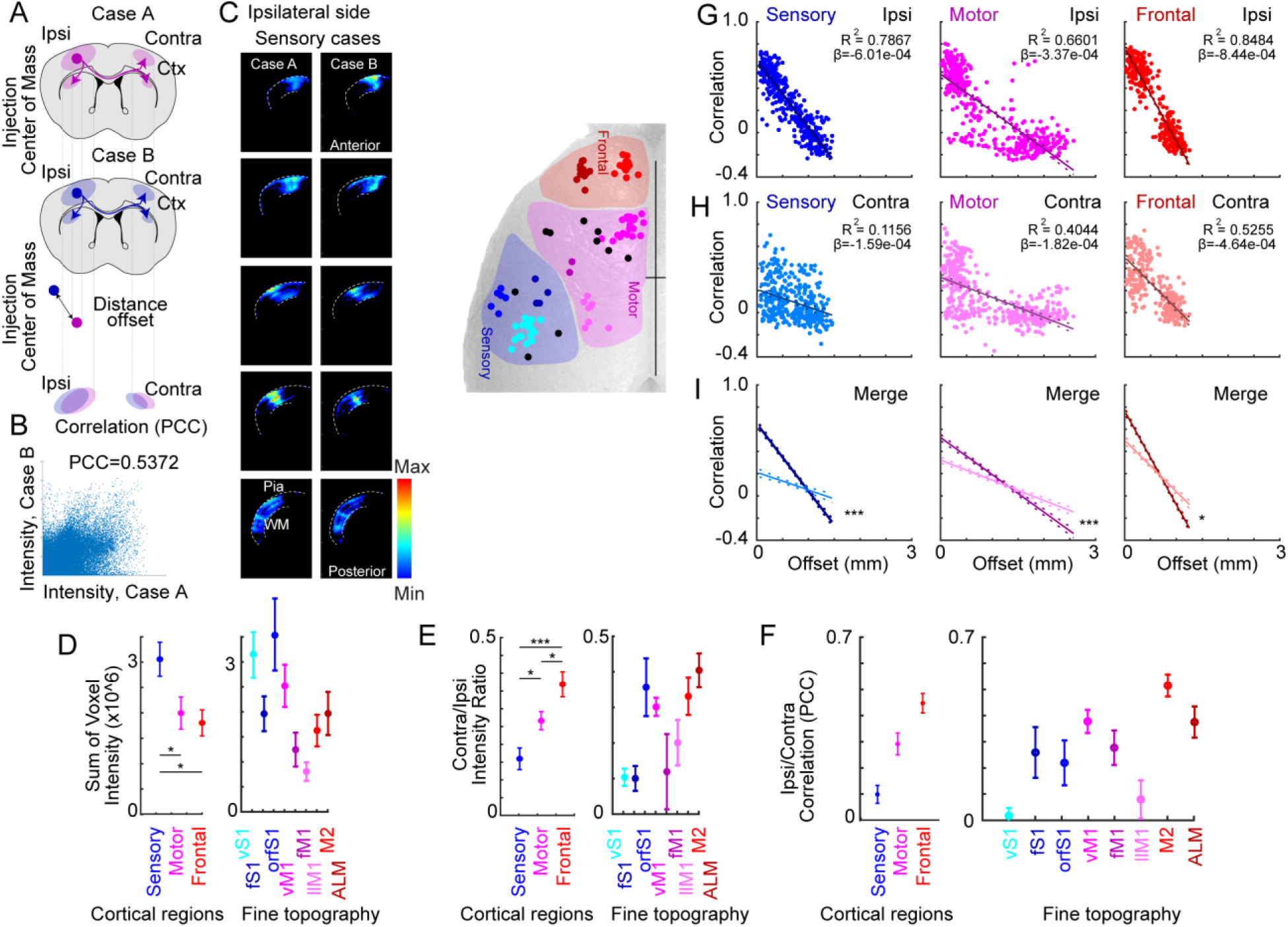
Topographic organization of contralateral cortical projections in frontal, motor, and somatosensory cortex. (A) Schematic illustrating how the corticocortical projections from two different injection sites are compared. The center of mass of cell bodies at the injection site is compared (offset distance in mm). The overlap of the ipsilateral of contralateral projections can be compared. (B) The correlation coefficient (PCC) for two example cases, comparing the suprathreshold voxel intensities for ipsilateral cortical projections. PCC=0.5372 for this example. (C) Sample coronal planes from two cases of aligned brains showing only suprathreshold ipsilateral cortical voxels. Spacing is ∼0.75 mm along the anterior/posterior axis, from ∼1.75 mm anterior to ∼1.25 mm posterior. Injection site locations in sensory, motor, and frontal locations as inset at right. (D) Measurement of the total voxel intensity for ipsilateral cortical projections. Voxels range from 0- 255 (in arbitrary units). Sensory injections were brighter than motor and frontal. Mean with SEM. 1 way ANOVA; df=2, F=4.89, p=9.97e-3. Tukey-corrected pairwise t-tests: Frontal–M1 (p=0.893), S1–M1 (p=0.035), S1–Frontal (p=0.015). Adjacent at right is the data subdivided for finer topographic divisions. (E) Ratio of voxel intensity for ipsilateral (injection site side) and contralateral voxels. 1 way ANOVA; df=2, F=12.07, p=2.66e-5. Tukey-corrected pairwise t-tests: Frontal–M1 (p=0.04), S1–M1 (p = 0.028), S1–Frontal (p = 1.4e-5). Adjacent at right is the data subdivided for finer topographic divisions. (F) Overall correlation between ipsilateral and contralateral corticocortical voxels by injection site, with subdivision of cortical sites into finer divisions at right. (G) Comparison of ipsilateral cortical correlations between injection sites for sensory, motor, and frontal injections. Mean with SEM. (H) Comparison of contralateral cortical correlations between injection sites for sensory, motor, and frontal injections. (I) Linear regression with 95% confidence interval. Two way ANCOVA examined the effects of region and ipsi/contra location on correlation (PCC) after controlling for distance (F(2,2165)=16.1, p=1.14e- 7). Pairwise comparisons were computed and Bonferroni adjusted and all regions had significantly higher slopes for ipsi versus contra: S1 (F(2165)=9.87, adj. p=1.63e-22), M1 (F(1,867) = 13.8, p=2.12e-4), and Frontal (F(2165)=2.33, adj. p=0.020).

In addition to making these comparisons across different injection cases, comparing each brain to the contralateral side allowed a direct assessment of symmetry. Correlations were computed for each injection site by comparing ipsilateral cortex to the aligned and reflected contralateral corticocortical projections (Fig. 4A). Example injections in aligned brains are shown for somatosensory, motor, and frontal (Fig. 4C-E) injections. As before, these sections of aligned brains are shown from anterior (top) to posterior (bottom). There is a remarkable difference in symmetry across the dorsal cortex. Frontal cortex injections have PCC ∼0.45, while motor areas are slightly symmetric <0.30 and somatosensory ones are very weakly symmetric at ∼0.10 (Fig. 4B). This is consistent with substantially greater symmetry in corticocortical innervation in anterior (as opposed to posterior) corticocortical projections.

**Figure 4.**
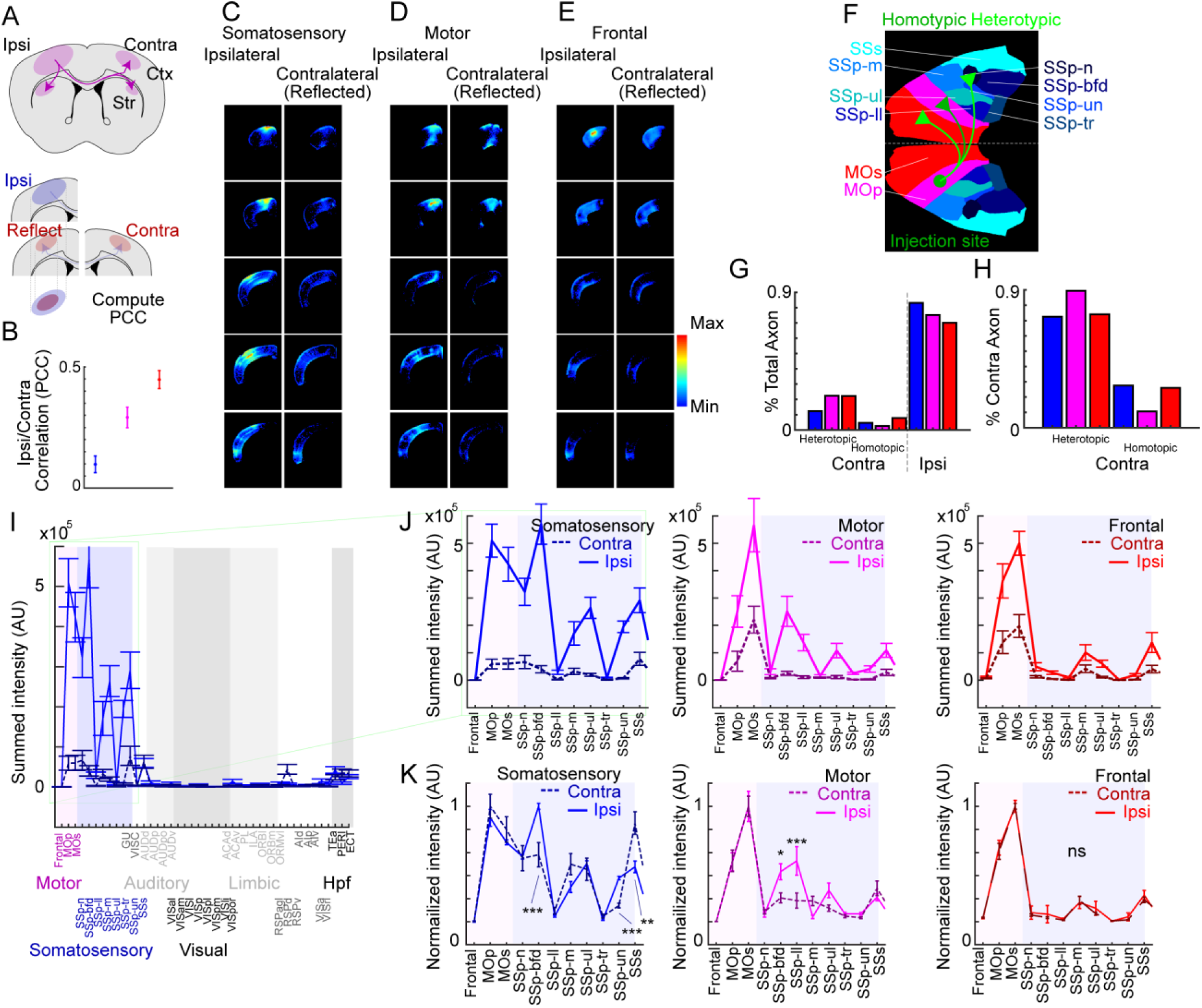
Area-by-area comparison of ipsi- and contralateral corticocortical projections in frontal, motor, and somatosensory cortex. (A) Schematic comparing corticocortical projections from a single injection site between ipsilateral and contralateral targets. (B) Average overall correlation (mean with SEM). (C-E). Sample coronal planes from aligned brains for somatosensory (C), motor (D), and frontal (E) injections. Left columns show ipsilateral cortex (injection side). Right columns show contralateral cortex, reflected for comparison. Only suprathreshold cortical voxels are shown. Spacing is ∼0.75 mm along the anterior/posterior axis, from +1.75 mm anterior to -1.25 mm posterior. (F) Topographic areas of cortex in the Allen CCFv3. To compare contralateral cortical projections, homotypic projections are contralateral projections from one cortical area (the injection site) to the corresponding cortical area (dark green), while heterotopic projections (light green) target contralateral cortex in areas not corresponding to the injection site. (G-H) Quantification of the fraction of contralateral corticocortical projections to ipsi- and contralateral areas based on suprathreshold voxel intensity. All somatosensory (blue), motor (purple), and frontal (red) injections are combined. (G) is normalized to all corticocortical projections. (H) is normalized to only contralateral projections. (I-K) Projection intensity to somatomotor cortical areas (per CCFv3) from somatosensory (blue), motor (purple), and frontal (red) injections. Regions are summed across cortical layers, noted on the x-axis in CCFv3 nomenclature. All areas are displayed for somatosensory cortex injections (I), with only somatomotor areas (J-K) displayed at right (where most axons are present). The summed intensity (in arbitrary units, AU) for suprathreshold voxels is plotted in (I) and (J). Solid lines are for ipsilateral projections and dotted lines are for contralateral injections. (K) represents peak normalized plots of the ipsilateral (solid) and contralateral (dotted) projection intensity. Ipsi- and contralateral intensity each normalized separately to compare the ipsi- and contralateral patterns in an intensity- independent manner. Error bars represent SEM. P-values represent ANOVA followed by posthoc Bonferroni-corrected t-tests.

Symmetry was further assessed as follows. First, corticocortical projections in a defined cortical region do not simply target the corresponding cortical area in the contralateral side of the brain, but also project to a range of ipsilateral and contralateral targets (Szczupak et al., 2023). For contralateral corticocortical output, homotypic projections are those that target the corresponding region of cortex in the other hemisphere. Heterotopic projections target other cortical areas, often adjacent ones (Fig. 4F). Using the CCFv3 regions to define these borders, IT-type contralateral axonal projections were indeed stronger to heterotopic than to homotopic targets (Fig. 4G-H), with heterotopic projections comprising 70-90% of contralateral projections. Because the CCFv3 contained several somatotopic subdivisions for somatosensory cortex, it was possible to assess the degree to which heterotopic axons targeted the same distribution of cortical areas in each hemisphere. Total axonal projection intensity to each CCFv3 cortical area (summed across layers) was computed and plotted across all cortical regions (shown for somatosensory cortex injections, Fig. 4I). Ipsilateral intensity (solid line) was higher than contralateral intensity (dotted line; ANOVA; df=2, F=24.55, p=2.37e-11). This was true for each cortical region individually as assessed with Bonferroni-corrected t-tests: S1 (p=4.8e-11), M1 (p=2.21e-11), and Frontal (p = 2.21e-6). Most projections were in somatomotor areas, so these were expanded in detail for sensory (blue), motor (purple), and frontal (red; Fig. 4J). Because contralateral intensity was generally lower, normalized distributions of ipsi- and contralateral intensity were plotted to compare the pattern (Fig. 4K). To compare normalized intensity across 48 cortical regions, a 3-way ANOVA was performed interacting cortical source region, cortical target region and ipsi/contra projection. All main effects and interactions were significant.

Therefore, we followed up with Bonferroni-corrected post-hoc t-tests. Differences in the main somatomotor areas are annotated in Fig. 4K: SSp-bfd, SSp-un, and SSs for somatosensory cortex and SSp-bfd and SSp-ll for motor cortex. Some other cortical target regions outside of somatomotor areas with weaker (not shown in Fig. 4K) did differ, including AId (for frontal projections), RSPv (for motor projections), and AIp, ECT, PERI, TEa, and VISC (for somatosensory areas). For somatosensory projections to medial temporal regions, these differences are consistent with the differences noted in Fig. 5C.

**Figure 5.**
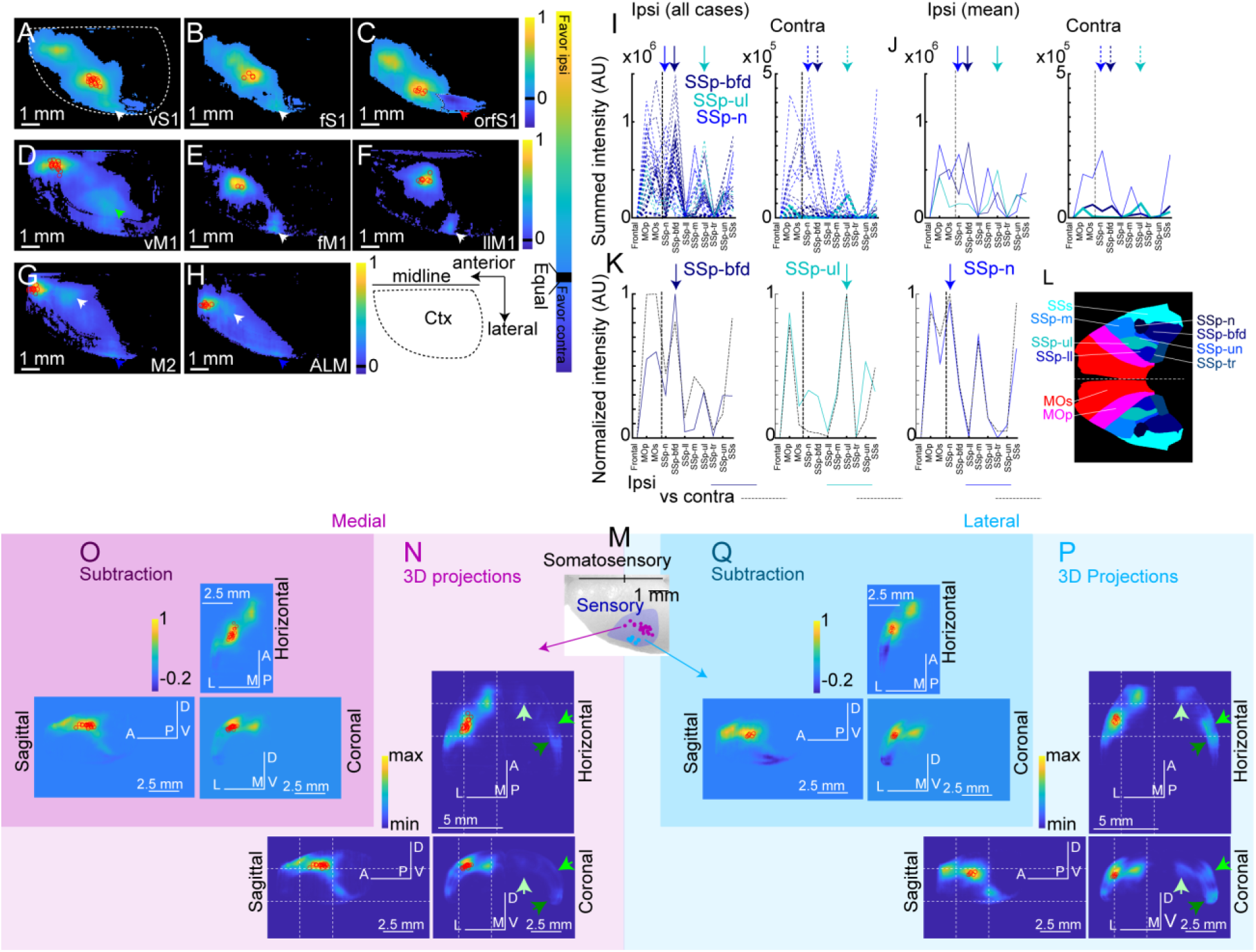
Symmetry of corticocortical projections in frontal, motor, and somatosensory cortex. (A-H) Injection sites in somatosensory (A-C), motor (D-F), and frontal (G-H) cortex, subdivided into topographic subregions (vS1, fS1, and orfS1 for somatosensory, A-C; vM1, fM1, llM1 for motor, D-F; and M2 and ALM for frontal, G-H). Injection sites are marked by red circles. Heatmap represents normalized projection intensity in ipsilateral cortex (summed across the dorsoventral axis), with contralateral intensity subtracted. Relative intensity is shown in the color map (yellow, stronger on the ipsilateral side; dark blue, stronger on the contralateral side). The outline of the dorsal surface of cortex is shown in (A). Black pixels represent values close to zero (contralateral and ipsilateral intensity roughly equal). (A-C) White arrowheads indicate strong ipsilateral connectivity with secondary somatosensory cortex (SSs). Red arrowheads indicate strong contralateral connectivity with entorhinal cortex. (D-F) Green arrowheads indicate strong motor projections to ipsilateral somatosensory cortex (SSp-bfd). White arrowheads indicate strong ipsilateral connectivity with secondary somatosensory cortex (SSs). (G-H) White arrowheads indicate strong ipsilateral connectivity with primary motor cortex (MOp). (I-K) Projection intensity to all cortical areas (per CCFv3) from subregions of somatosensory cortex injections. Left plot (I and J) shows the summed intensity (in arbitrary units, AU) for suprathreshold voxels in ipsilateral cortex for each individual case (color coded to the subregion), with arrow at top indicating the CCFv3 region best corresponding to the injection site. The right plot shows the same data for the contralateral side. The mean of the ipsilateral (left) and contralateral (right) is presented in (J). Normalized plots of ipsilateral (blue colors) and contralateral (black) are presented in (K) for vS1 (left), fS1 (middle), and orfS1 (right). (L) Color coded plot of the CCFv3 areas for somatosensory cortex. (M) Injection sites in somatosensory cortex divided into more medial (N=21, violet) and lateral (N=6, azure) categories. (N) The cortical voxels from both left and right cortex were peak normalized for each medial injection case, then averaged and renormalized. The resulting cortical voxels were projected into horizontal (top), sagittal (left), and coronal projections to show axon targeting. Red circles indicate center of mass of the injection sites. Orientation of panels is indicated with A/P (anterior/posterior), M/L (medial/lateral), and D/V (dorsal/ventral). The white dashed lines indicate the same planes in both medial and lateral images for comparison. Bright, middle, and dark green arrows indicate location of contralateral M1, S1, and S2, respectively, for comparison. (O) The contralateral voxels were reflected and subtracted from ipsilateral voxels to present a difference map. (P, Q) Data presented in a similar fashion for lateral S1 injections.

Because of the alignment to a standard atlas, symmetry was assessed by reflecting the IT-type projections from the contralateral side and subtracting them from the ipsilateral side (as was done above to assess homotypic projections). Because the structures are 3D, they are presented in figures as a projection (summing along the dorsoventral axis). For cortex, this results in a map that appears as a view from the dorsal surface (Fig. 5A-H). Injection site intensity (marked with red circles) is reflected in yellow with high ipsilateral intensity. Cortical areas unrelated to somatomotor injection sites are close to background (appearing black).

These maps helped identify areas with substantial ipsilateral projections, such as connections between M1 and S1, as well as projections to entorhinal cortex. In S1 injections, corticocortical projections target secondary somatosensory cortex (S2) as well as entorhinal areas in both ipsi- and contralateral hemispheres. Remarkably, contralateral entorhinal cortex received stronger projections from orfS1(white dotted line and red arrow in Fig. 5C) (Mao et al., 2011). Of note, there are differences between more medial and lateral injection sites in S1, with lateral S1 regions showing stronger callosal projections (Montanari et al., 2023) to M1, S1 and S2 (Fig. 5M-Q). Motor areas send projections to principally ipsilateral somatosensory cortex, though this is most apparent in vM1 injections (green arrow, Fig. 5D), as whisker areas in M1 and S1 are sufficiently separate to distinguish projections. Forelimb M1 and S1 are adjacent, which makes this more difficult to test (but ipsilateral sites appear interconnected in Fig. 5B). These areas also project to entorhinal areas (white arrows, Fig. 5E-F). Frontal areas also have strong connectivity with ipsilateral motor areas (white arrows, Fig. 5G-H). Because of the somatosensory subdivisions within CCFv3, it was also possible to individually assess vS1 (SSp- bfd), fS1 (SSp-ul), and orfS1 (SSp-n) in the same manner (Fig. 5I-L). These largely showed different distributions between different areas (vS1 versus fS1) but similar patterns between ipsilateral and contralateral projections (Fig. 5K).

### High correlation of projection targeting in IT-type corticostriatal projections

Similar patterns were present in a subset of corticostriatal projections examined (Fig. 6). Corticostriatal projections target topographically defined regions of striatum (Foster et al., 2021; Hintiryan et al., 2016; Hooks et al., 2018). Thus, frontal cortex densely targets large areas of striatum, including portions of ventral and dorsolateral striatum (Fig. 6A-C), while motor (Fig. 6D-F) and somatosensory (Fig. 6G-I) cortices target mainly dorsolateral striatum. To compare the ipsi- and contralateral projections to striatum (Fig. 6L-N) at different rostrocaudal planes, images from injection sites in frontal cortex (Fig. J-K) are shown, with detail in the striatum at three rostrocaudal planes (Fig. 6O-T).

**Figure 6.**
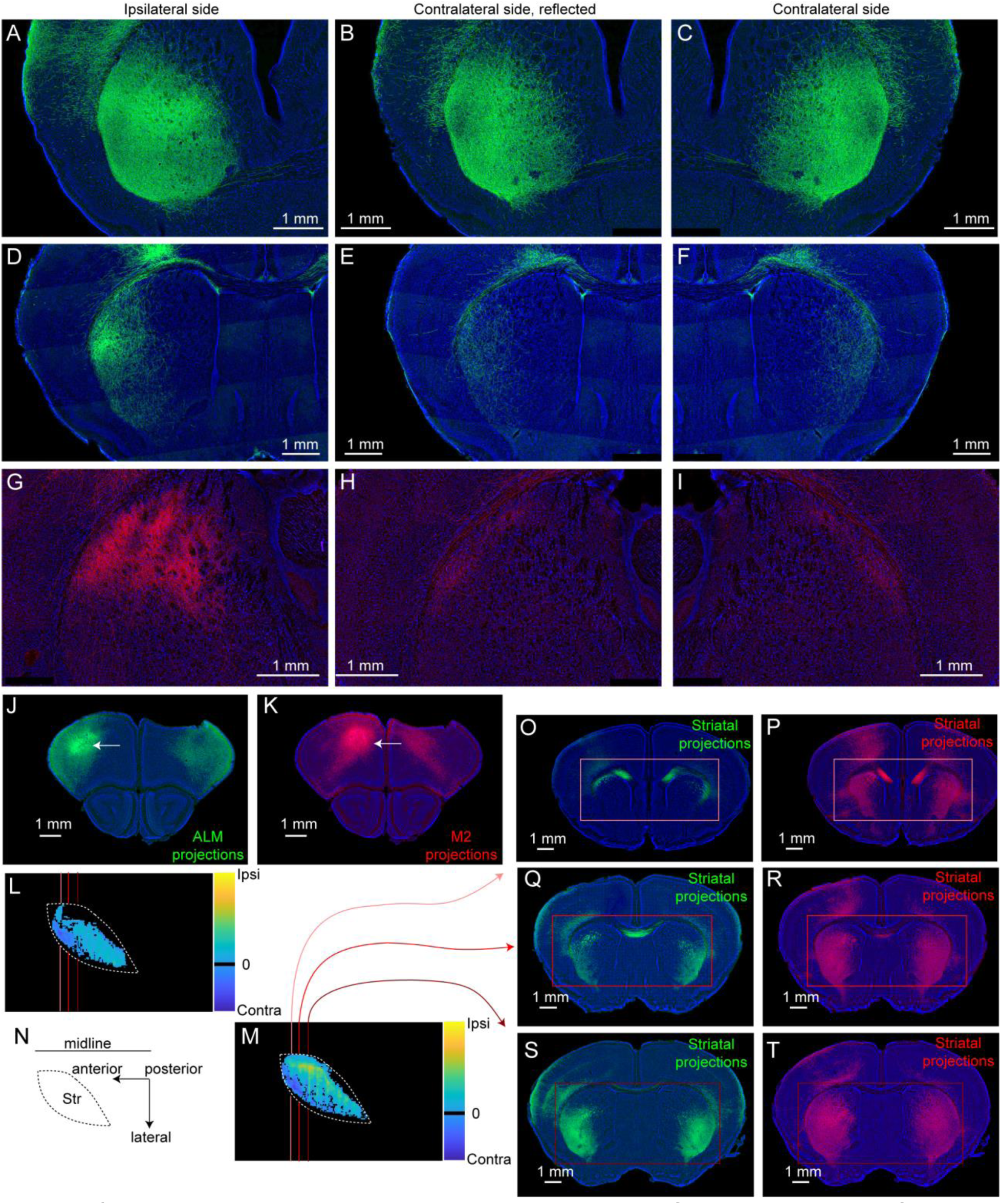
Ipsi- and contralateral corticostriatal projections of IT-type neurons in frontal, motor, and somatosensory cortex. (A-I) Coronal images showing injection site of AAV-DIO-XFP into frontal cortex (A-C), motor cotex (D-F), and somatosensory cortex (G-I). Left column shows ipsilateral side, right column shows contralateral side, with reflected image shown for comparison. (J-T) More detailed presentation of the injection sites in ALM (J, green) and M2 (K, red) and their striatal targets. Arrowheads show axonal projections. (L-N) present the overall difference in striatal axonal intensity between ipsilateral and contralateral projections (see Fig. 8) for the ALM (L) and M2 (M) injections, with a schematic of the dorsal view of striatum (Str). Red lines indicate approximate planes of the images in (O-T), with the striatum boxed in the corresponding color.

Injections were quantitatively compared as before. The distance offset was calculated in millimeters, while the correlation coefficient of overlapping suprathreshold voxels was computed for pairs of injections (Fig. 7A-B). Striatal voxel identity was determined by registration to the Allen CCFv3. Example images of the contralateral striatum are shown for two aligned brains for a frontal injection site, with planes spaced at 0.75 mm ranging from ∼1.50 mm anterior to bregma to ∼1.50 mm posterior (Fig. 7C). Striatal projections were strongest from frontal areas in mouse (Fig. 7D), with summed voxel intensity more than twice that of motor and somatosensory injections. Furthermore, projections to contralateral striatum were also relatively brighter in frontal injections based on the ratio of contralateral/ipsilateral intensity (Fig. 7E). This effect was more strongly pronounced in striatum than in cortex.

As for cortical injections, injection sites in nearby locations in aligned brains showed high degrees of correlation in the targeting of their ipsilateral striatal projections (Fig. 7G), which has previously been examined in detail (Hooks et al., 2018). Adding to this prior study of motor and sensory corticostriatal topography, the frontal injections also show high topographic organization, with the ipsilateral corticostriatal projections of frontal cortex (red) being perhaps more precise than somatosensory ones. Peak PCC with no offset is ∼0.7 (linear fit near 0 offset). In comparison, peak somatosensory PCC (royal blue) is ∼0.6 and motor (purple) is ∼0.4. Thus, ipsilateral corticostriatal axons also reliably target to similar cortical areas across different animals. Contralateral striatal projections also showed an unexpected pattern. As in cortex, these showed correlation that varied with injection site offset (Fig. 7H). But somatosensory cortex, which had remarkably high correlation in ipsilateral connectivity, had essentially no correlation (light blue) on the contralateral side (Fig. 7H). Further, contralateral motor cortex projections to striatum (light purple) were not as well organized as ipsilateral projections. As in cortex, there were not as weak as somatosensory ones. But contralateral corticostriatal projections from frontal cortex (pink) were remarkably similar. The fit was difficult to distinguish between ipsi- and contralateral projections (Fig. 7I).

**Figure 7.**
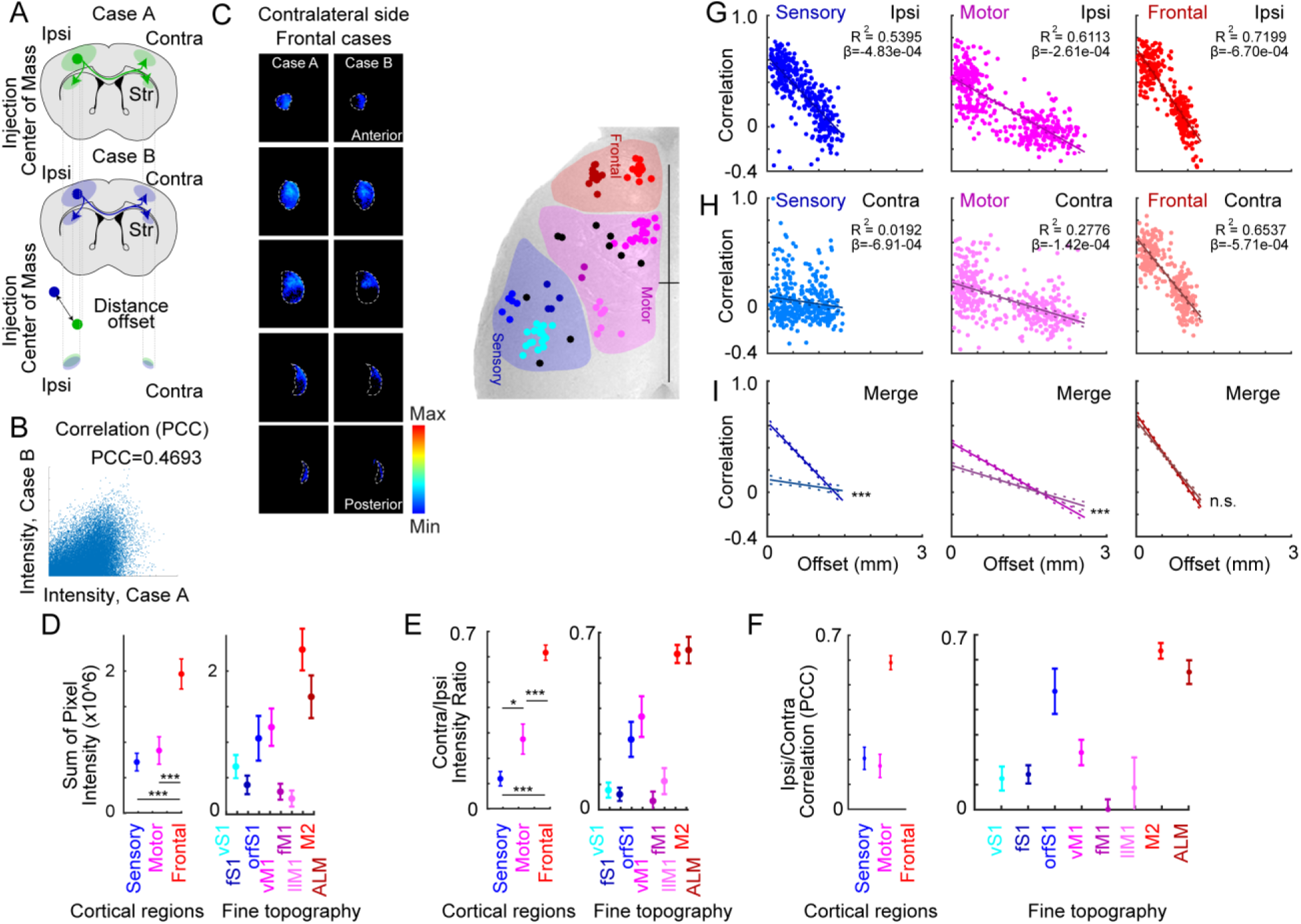
Topographic organization of contralateral corticostriatal projections in frontal, motor, and somatosensory cortex. (A) Schematic illustrating how the corticostriatal projections from two different injection sites are compared. The center of mass of cell bodies at the injection site is compared (offset distance in mm). The overlap of the ipsilateral of contralateral projections can be compared. (B) The correlation coefficient (PCC) for two example cases, comparing the suprathreshold voxel intensities for ipsilateral striatal projections. PCC=0.4693 for this frontal example. (C) Sample coronal planes from two cases of aligned brains showing only suprathreshold contralateral striatal voxels. Spacing is ∼0.75 mm along the anterior/posterior axis, from ∼1.50 mm anterior to ∼1.50 mm posterior. Injection site locations in sensory, motor, and frontal locations as inset at right. (D) Measurement of the total voxel intensity for ipsilateral striatal projections. Voxels range from 0-255 (in arbitrary units). Frontal injections were brighter than somatosensory and motor. Mean with SEM. 1 way ANOVA; df=2, F=13.63, p=8.21e-6. Tukey-corrected pairwise t-tests: Frontal–M1 (p=1.5e-4), S1–M1 (p=0.787), S1–Frontal (p=2.1e-5). Adjacent at right is the data subdivided for finer topographic divisions. (E) Ratio of voxel intensity for ipsilateral (injection site side) and contralateral voxels. 1 way ANOVA; df=2, F=32.84, p=4.18e-11. Tukey-corrected pairwise t-tests: Frontal–M1 (p=1.0e-6), S1–M1 (p=0.03), S1–Frontal (p<1e-16). Adjacent at right is the data subdivided for finer topographic divisions. (F) Overall correlation between ipsilateral and contralateral corticostriatal voxels by injection site, with subdivision of cortical sites into finer divisions at right. (G) Comparison of ipsilateral striatal correlations between injection sites for sensory, motor, and frontal injections. (H) Comparison of contralateral striatal correlations between injection sites for sensory, motor, and frontal injections. (I) Linear regression with 95% confidence interval. Two way ANCOVA examined the effects of region and ipsi/contra location on correlation (PCC) after controlling for distance (F(2,2165)= 67,565, p=3.45e-29). Pairwise comparisons were computed and Bonferroni adjusted and all regions had significantly higher slopes for ipsi versus contra: S1 (F(2162)=16.7, adj. p=3.46e-59), M1 (F(2162)=6.46, adj. p=1.3e-10), and Frontal (F(2162)=2.33, adj. p=0.91).

This suggested a high degree of symmetry in the strong corticostriatal projections. Thus, the contra- and ipsilateral side of each brain were compared by reflecting the registered contralateral projections (Fig. 8C-D). Examples are shown for both motor (Fig. 8C) and frontal cases (Fig. 8D), with frontal projections having much stronger connectivity to ventral striatum.

**Figure 8.**
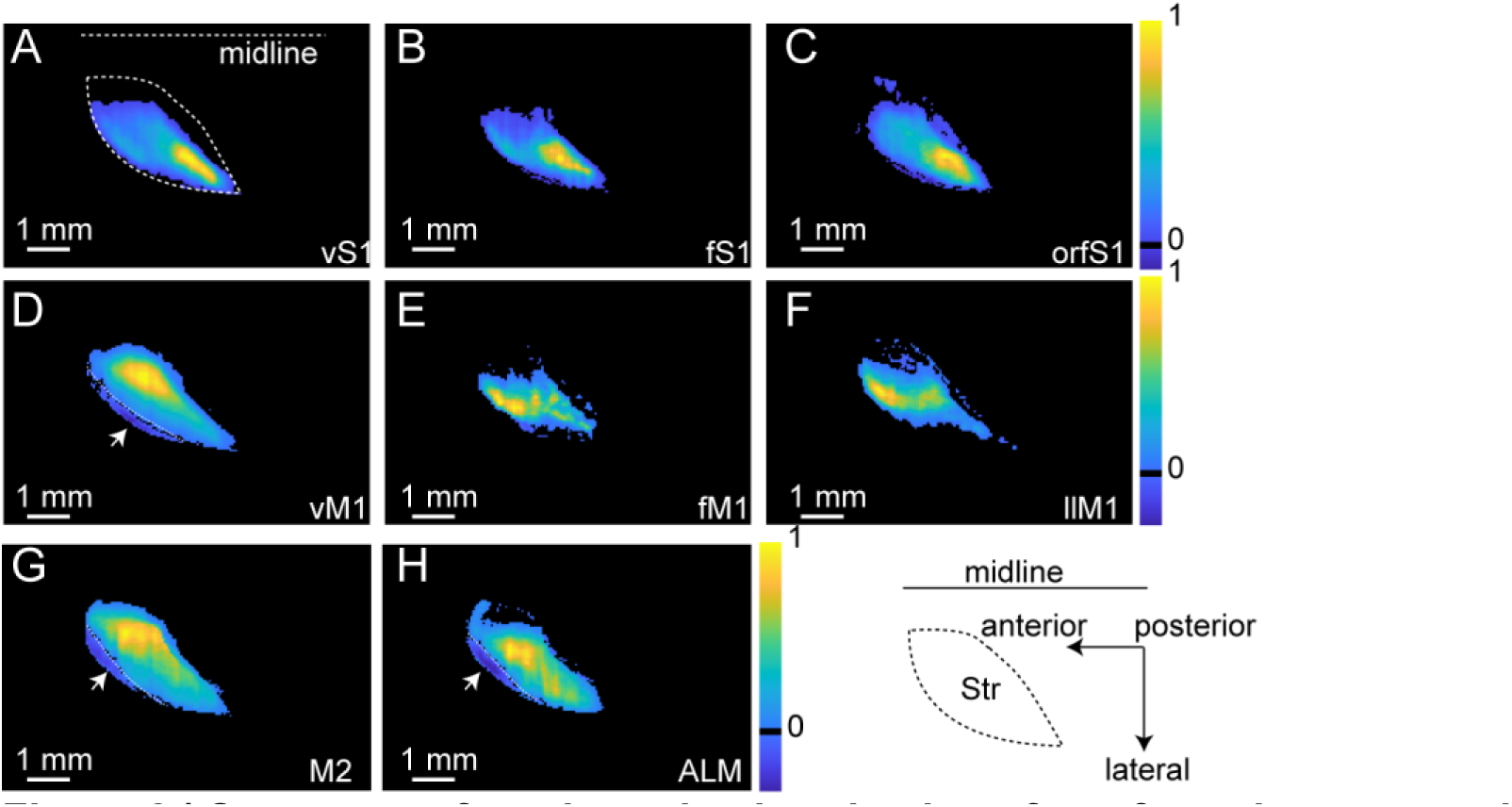
Symmetry of corticostriatal projections from frontal, motor, and somatosensory cortex. (A-H) Striatal projections from injection sites in somatosensory (A-C), motor (D-F), and frontal (G-H) cortex, subdivided into topographic subregions (vS1, fS1, and orfS1 for somatosensory, A-C; vM1, fM1, llM1 for motor, D-F; and M2 and ALM for frontal, G-H). Injection sites are not marked. Heatmap represents normalized projection intensity in ipsilateral striatum (summed across the dorsoventral axis), with contralateral intensity subtracted. Black pixels represent values close to zero (contralateral and ipsilateral intensity roughly equal), with yellow representing sites with stronger ipsilateral strength (injection sites) and dark blue voxels representing stronger contralateral sites. White arrowheads reflect anterolateral sites of strong contralateral connectivity. Schematic inset at bottom right shows the shape of striatum as viewed from the dorsal side, with midline and anterior/posterior axis labeled.

Symmetry was impressive for frontal projections, with the average PCC ∼0.6 for frontal cortex. In marked contrast, this symmetry was largely absent in motor and somatosensory cases, with PCC ∼0.2 for both. Having established that there was a rostrocaudal gradient in the symmetry of corticostriatal projections, it was of interest to test for a medial-lateral gradient. The frontal cortex injections used could readily be sorted into more medial (M2) or lateral (ALM) locations (Hooks et al., 2018) (note the names for these regions vary; see Discussion). Comparing these did not reveal substantial differences in the degree of symmetry between frontal injection sites (Fig. 7F). Overall, this data suggests that corticostriatal projections can show even greater symmetry than corticocortical projections, and that this symmetry varies principally along the rostrocaudal axis, with symmetry highest in frontal areas. Note that this was not tested in posterior parietal areas or other sensory areas such as auditory and visual cortex.

Taking advantage of the alignment to a standard atlas, symmetry was similarly assessed by reflecting the projections from the contralateral striatum and subtracting from the ipsilateral striatum. Striatal voxels are then presented as a dorsoventral projection, summed along the dorsoventral axis (Fig. 8A-H). For striatum, this results in a paisley-shaped map viewed from above. High projection site intensity is reflected in yellow for high ipsilateral intensity. Unrelated striatal areas are close to background (black). These maps were largely consistent with the topographic targeting of somatosensory and frontal corticostriatal projections previously described, with slight differences between cortical injection site subdivisions (Fig. 8A-F).

However, this map also identified areas in the anterolateral striatum (arrowheads and dotted lines) where contralateral corticostriatal projections were more intense than ipsilateral ones (Fig. 8D,G,H). This might be one small specialization in the contralateral corticostriatal projection.

Overall, these results suggest largely symmetrical corticostriatal projections in frontal areas, which are also stronger than those in motor or somatosensory cortex.

### Weaker innervation and low correlation of contralateral PT-type and CT-type corticothalamic projections

Cre-driver mouse lines selective for PT-type (Fig. 9, 10, and 13) and CT-type (Fig. 11, 12, and 14) neurons were injected with AAV expressing Cre-dependent tracers, including GFP, td-tomato, and smFPs. The PT-type mouse lines show excellent laminar restriction to L5B (which houses most PT-type neurons) in all cortical areas studied (Fig. 9A-E). Whole brain imaging and reconstruction confirmed that PT-type neurons project ipsilaterally to striatum, and then target downstream areas including thalamus and brainstem. These projections mainly arborize in ipsilateral thalamus (Fig. 10), though some axons cross the midline. To visualize thalamic targets, we constructed 3D models of thalamic innervation using aligned brains from each injection. Only suprathreshold voxels in the thalamus were included, and each injection was normalized to its peak. Frontal, motor, and sensory injections were considered together, with an average thalamic map for each of these three areas generated by averaging normalized cases. These are displayed in horizontal, sagittal, and coronal views for the ipsilateral (Fig. 13A) and contralalateral thalamus (Fig. 13B). For these PT-type corticothalamic projections, ipsilateral innervation is substantially denser, and the ratio significantly favors the ipsilateral side (Fig. 13D). This is most pronounced for sensory regions, whose main thalamic targets include the higher order thalamic nucleus PO, which is relatively lateral (Veinante et al., 2000). Motor areas project to more medial thalamic nuclei, such as VAL, and VM, though these still include PO (Guo et al., 2020; Hooks et al., 2013). Frontal areas show much weaker L5B corticothalamic afferentation and very weak contralateral afferents as well. This results in very poor correlation of ipsi- and contralateral targeting (Fig. 13E) and difference plots that reflect the distribution of the stronger ipsilateral projections, especially for somatosensory projections.

**Figure 9.**
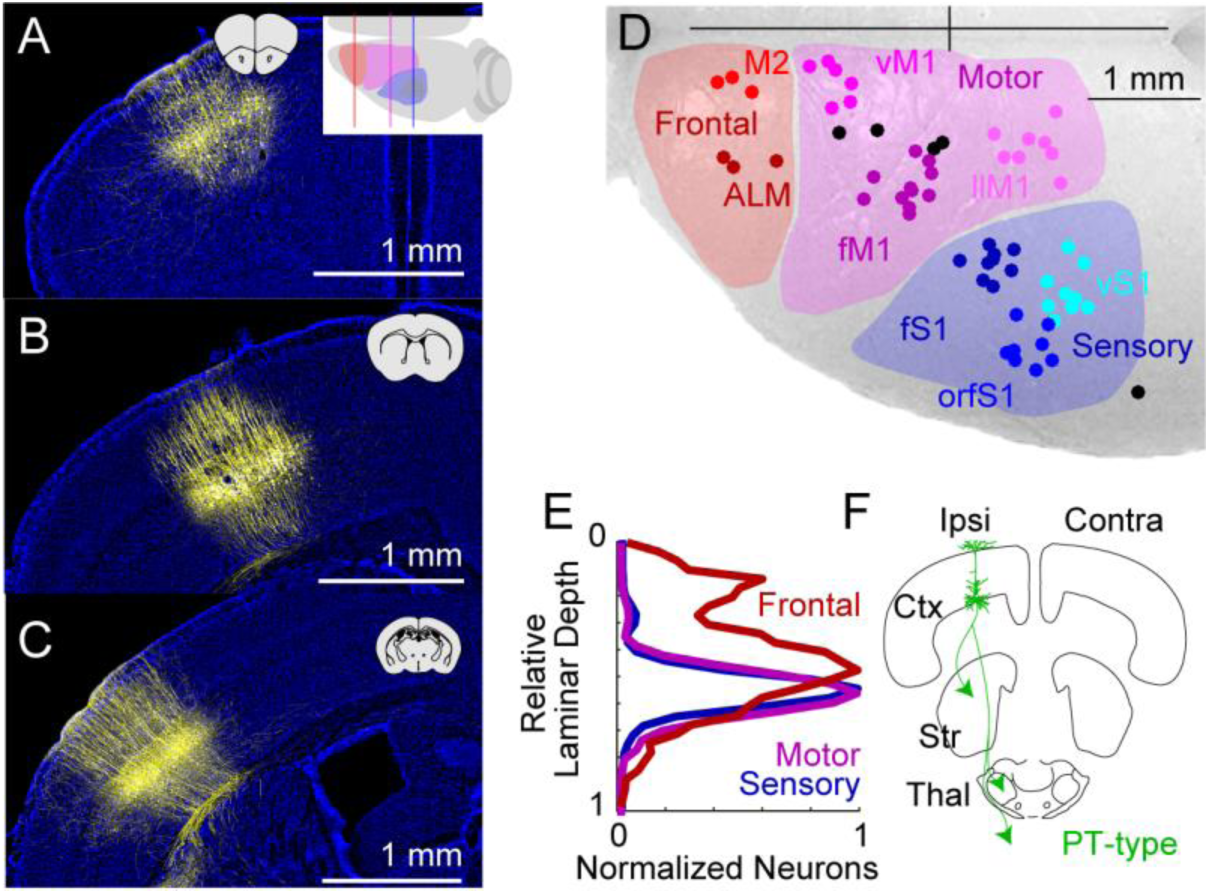
Mapping layer 5 corticothalamic output from sensory and motor cortex. Injection site of AAV-DIO-XFP into frontal (A), motor (B), and sensory cortex (C) of Sim1_KJ18-Cre+ mice. Inset of coronal brain slices shows approximate location. In (A), top-down cartoon shows approximate planes of section. (D) Center of mass of all injection sites in sensory (blue, N=26), motor (purple, N=25), and frontal (red, N=6) cortex. Subdivisions within areas (M2, ALM, vM1, fM1, llM1, vS1, fS1, and orfS1) are indicated in different colors, with uncategorized injection sites in black. Bregma is indicated as the cross at top right. (E) Quantification of relative depth of labeled Cre+ cortical neurons in frontal (red), motor (purple), and sensory (blue) cortex. (F) Cartoon of output targets of PT-type neurons, including striatum (Str), thalamus (Thal), and brainstem.

**Figure 10.**
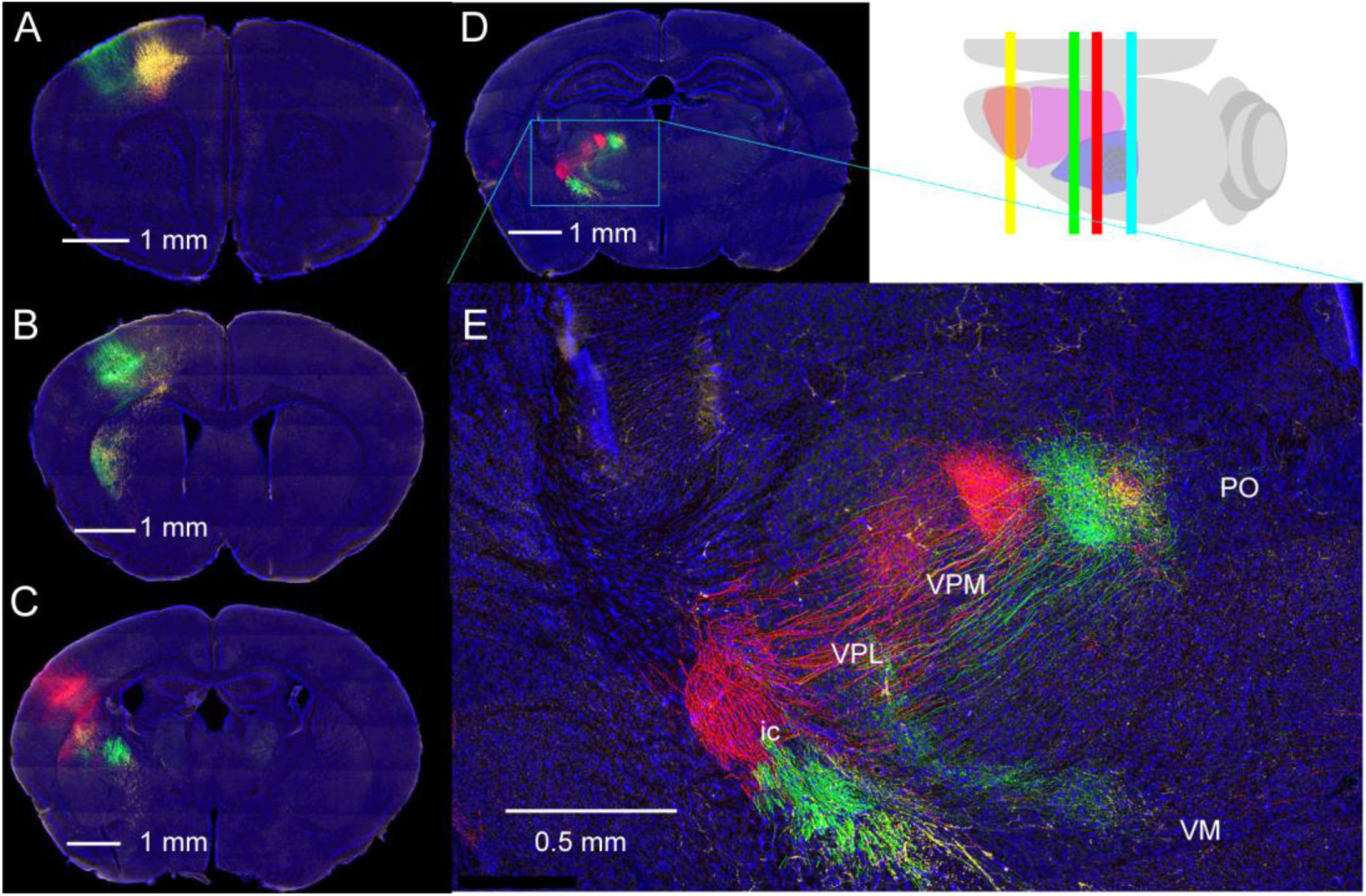
Ipsilateral corticothalamic projections of PT-type neurons in frontal, motor, and somatosensory cortex. (A-C) Coronal images showing injection site of AAV-DIO-XFP into frontal (A), motor (B), and somatosensory (C) cortex, as indicated in the cartoon at top right. (D) Example plane of thalamic projections to posterior thalamus (PO) shown. Note the absence of contralateral projections. (E) Axons in internal capsule (ic) shown, with some collaterals in ventrobasal thalamus (VB, comprised of VPM and VPL) and axons projecting towards ventromedial thalamus (VM).

**Figure 11.**
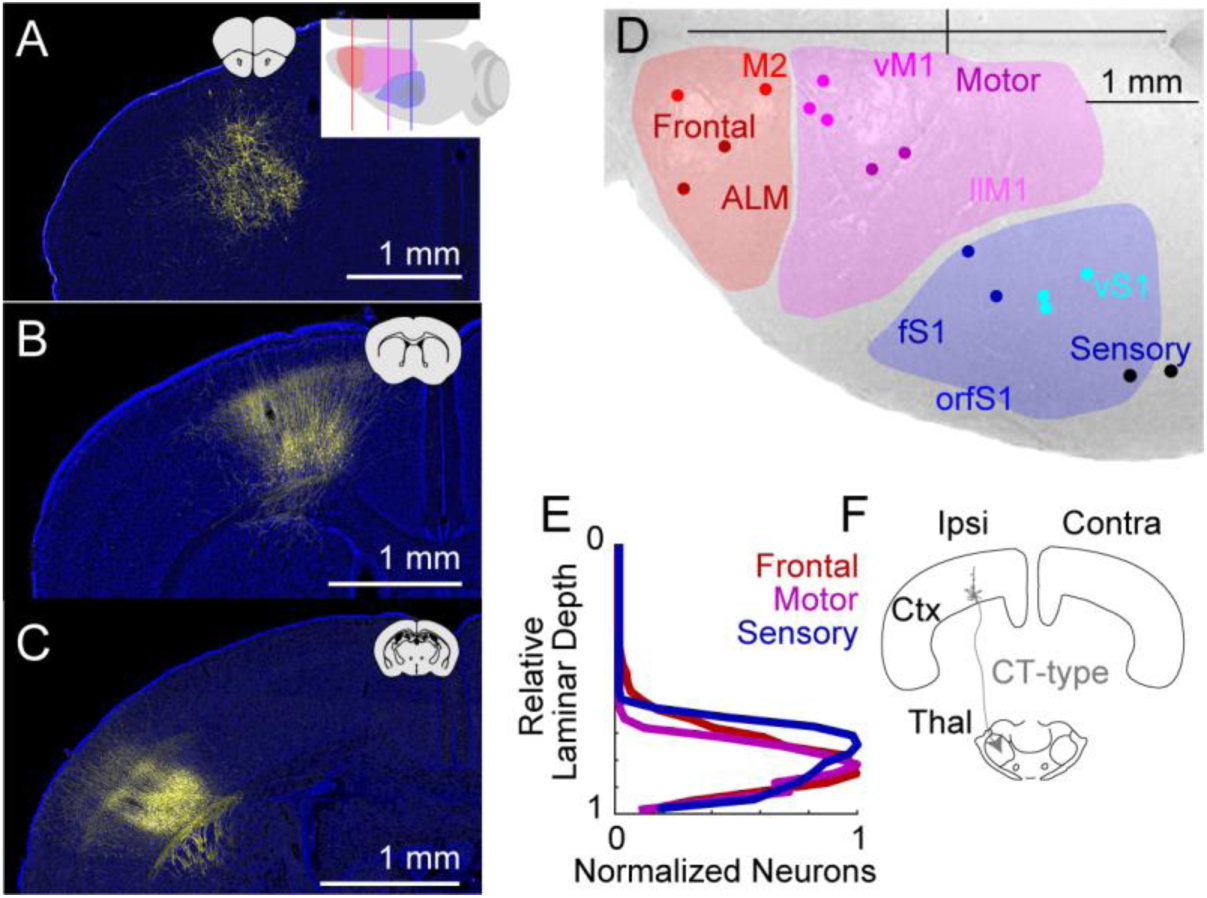
Mapping layer 6 corticothalamic output. Injection site of AAV-DIO-XFP into frontal (A), motor (B), and sensory cortex (C) of Ntsr1_GN220-Cre+ mice. Inset of coronal brain slices shows approximate location. In (A), top-down cartoon shows approximate planes of section. (D) Center of mass of all injection sites in sensory (blue, N=5), motor (purple, N=5), and frontal (red, N=4) cortex. Subdivisions within areas (M2, ALM, vM1, fM1, vS1, and fS1) are indicated in different colors, with uncategorized injection sites in black. Bregma is indicated as the cross at top right. (E) Quantification of relative depth of labeled Cre+ cortical neurons in frontal (red), motor (purple), and sensory (blue) cortex. (F) Cartoon of output targets of CT-type neurons, including thalamus (Thal).

**Figure 12.**
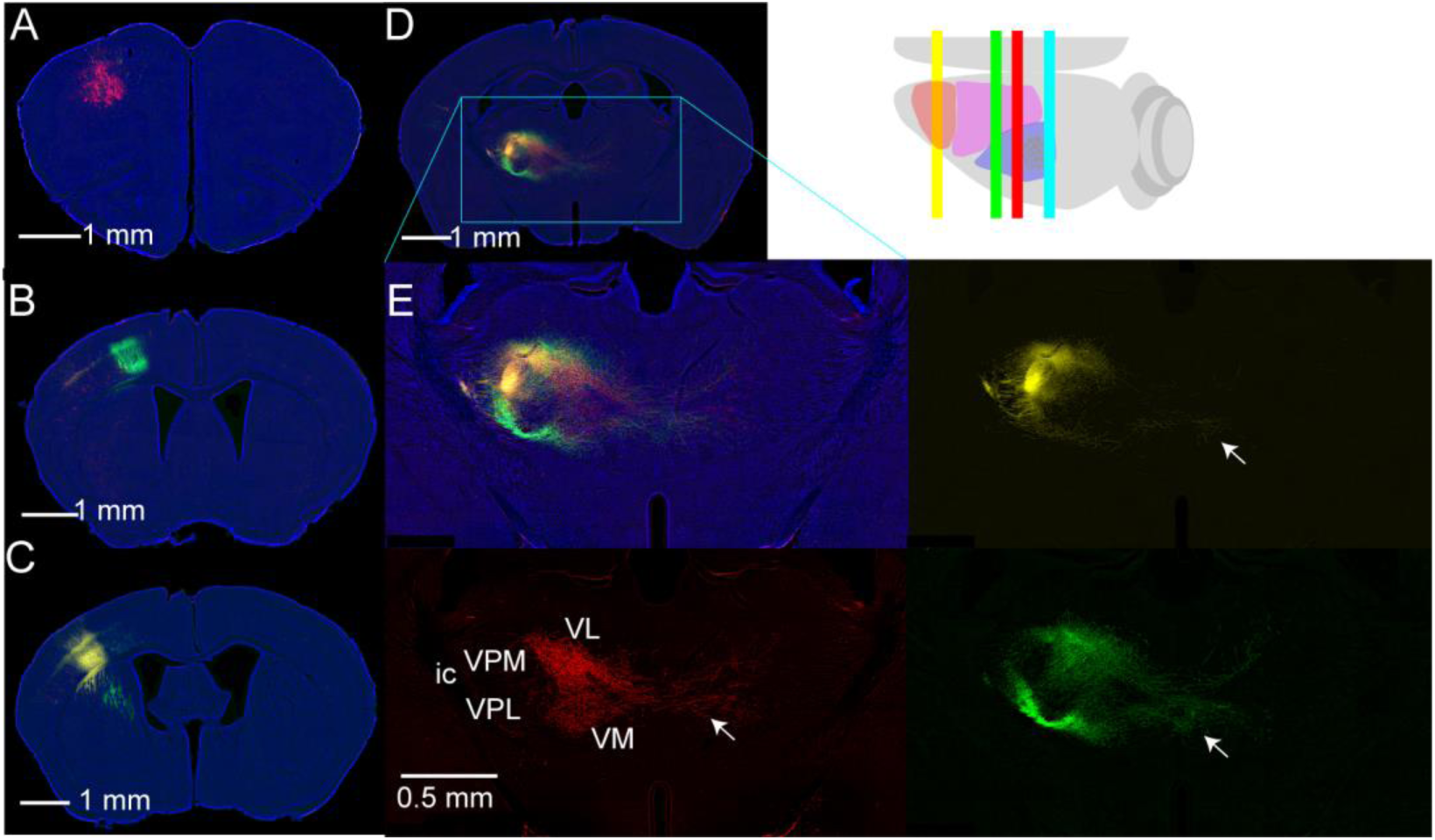
Corticothalamic projections of CT-type neurons in frontal, motor, and somatosensory cortex. (A-C) Coronal images showing injection site of AAV-DIO-XFP into frontal (A), motor (B), and somatosensory (C) cortex, as indicated in the cartoon at top right. (D) Example plane of thalamic projections to posterior thalamus (PO) shown. Note the absence of contralateral projections. (E) Axons in internal capsule (ic) shown, with some collaterals in ventrobasal thalamus (VB, comprised of VPM and VPL) and axons projecting towards ventrolateral (VL) and ventromedial thalamus (VM).

**Figure 13.**
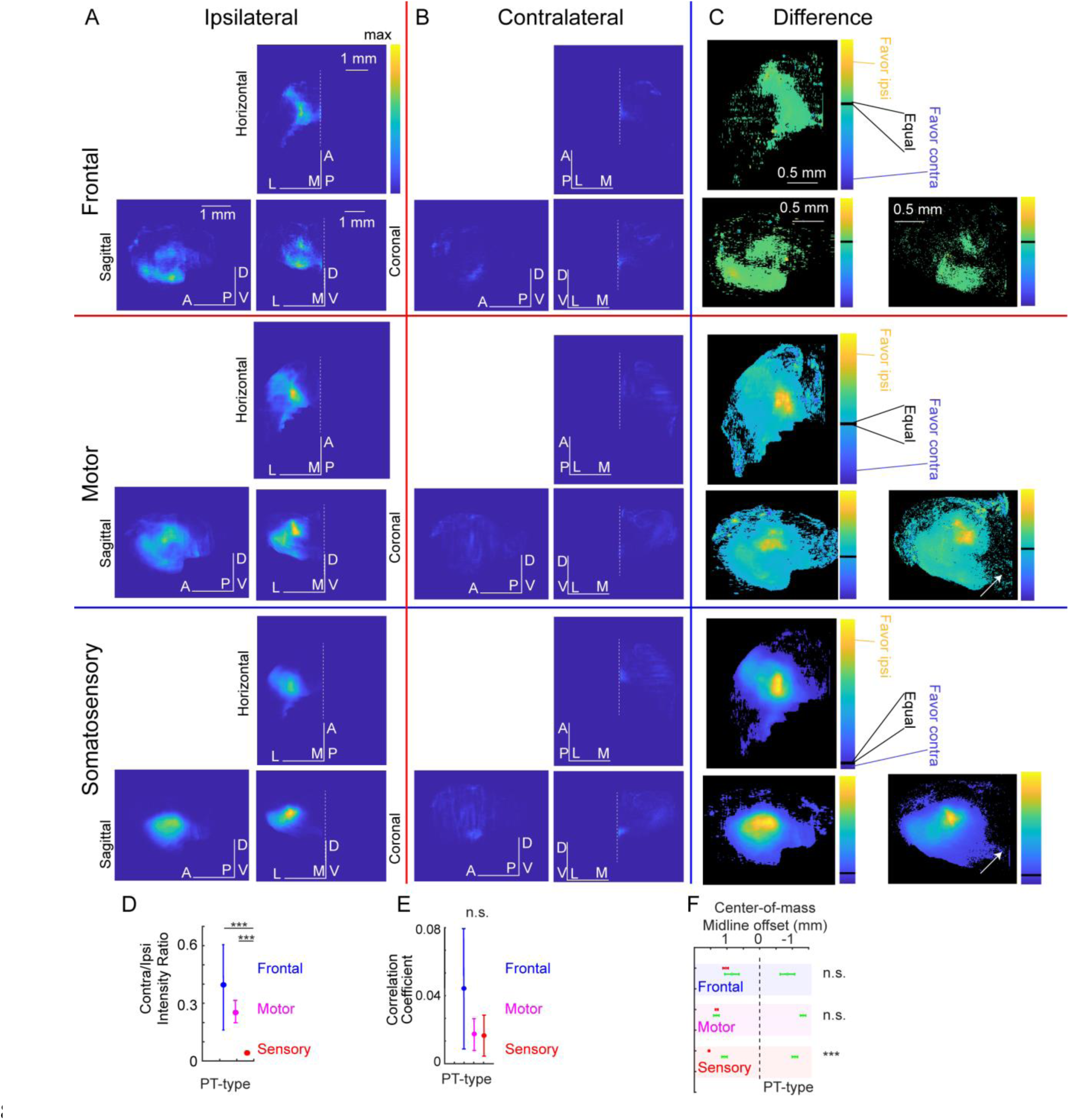
Comparison of ipsi- and contralateral PT-type corticothalamic projections across frontal, motor, and somatosensory cortex. (A) A 3D map of PT-type projections to the thalamus was generated from each injection by selecting only thalamic voxels and nomalized to the peak. These were then averaged with similar injection sites (frontal injections in the top row, motor injections in the second row, and somatosensory injections in the last row). The averaged normalized voxels for the left (ipsilateral) injection site were then displayed as projections along the horizontal (top), sagittal (left) or coronal plane (right). Anterior/posterior (A/P), dorsal/ventral (D/V), and medial/lateral (M/L) as indicated. Midline is shown as a dotted white line in horizontal and coronal planes. (B) The same presentation for the contralateral projections. (C) The difference between ipsilateral and contralateral was computed by subtraction for each brain, then averaged. The subtraction is shown as the left (ipsilateral) view, with black indicating values closest to zero. Arrows in the motor and sensory rows show comparable input where midlnie crossing axons are present. (D) Ratio of intensity of contralateral and ipsilateral projections shown as the mean +/- SE for frontal (blue), motor (purple), and somatosensory (red). (E) Correlation coefficient for ipsi- and contralateral thalamus (note y-axis compared to scores for striatum). (F) Center of mass of ipsilateral (red, left) and contralateral (green, right) thalamic voxels plotted as mean +/- SE from the midline (black dashed line). Contralalateral position plotted in a mirrored location at left for statistical comparison to ipsilateral midline offset. t-test used for all statistical comparions. *p<0.05; **p<0.01; ****; p<0.001.

The corticothalamic projections of L5B PT-type and L6 CT-type neurons have long been known to have differences in anatomical appearance, targeting, and synaptic function, with the former being referred to as driver synapses and the later as modulatory (Sherman & Guillery, 2006). It is worth noting, however, that there are vastly more L6 neurons than L5B (Keller et al., 2018). We repeated this experiment in Cre-driver mouse lines selective CT-type (Fig. 11, 12, and 14) neurons. The CT-type mouse line had good laminar restriction to L6 (Fig. 11A-E). CT- type neurons project to thalamus with limited collateralization in cortex. As with PT-type neurons, these projections mainly arborize in ipsilateral thalamus, though midline-crossing axons are visible (Fig. 12E).

The average thalamic map for CT-type projections in each of these three areas was generated by averaging normalized cases as before. Maps were more robust in part due to the denser and brighter thalamic innervation. These are displayed in horizontal, sagittal, and coronal views for the ipsilateral (Fig. 14A) and contralalateral thalamus (Fig. 14B). For CT-type corticothalamic projections, contralateral innervation, though less dense, was now sufficient to readily visualize differences in ipsi- and contralateral targeting. The brightness of these inputs still favors the ipsilateral side, most strongly for sensory and most weakly for frontal areas (Fig. 14D). The main thalamic targets differ slightly from PT-type inputs, with somatosensory targets preferring VPM and VPL (the principal somatosensory thalamus) to PO (higher order thalamus) (Sherman & Guillery, 2006; T. Kim, 2020). This results in strong, focal targeting of VPM/VPL in sensory projections. The difference plots show relatively similar (black) levels of innervation of midline thalamic nuclei, including the reuniens and rhomboid nuclei (arrows in Fig. 12E and yellow axons in Fig. 14C). However, overall correlation between ipsi- and contralateral projections are poor (<0.1, Fig. 14E) because of the general strength of ipsilateral projections vs. contralateral ones. Though contralateral projections were relatively stronger in motor areas, ipsilateral ones were still predominant, targeting more dorsal/lateral areas of VL and PO, with some overlap in the midline nuclei (reuniens and rhomboid nuclei; arrows in Fig. 12E and green axons in Fig. 14C). Motor areas did also project to medial motor thalamic nuclei, including ipsilateral VM. Frontal areas had weaker CT-type corticothalamic input but sufficient to contrast between the hemispheres. Correlation was poor between ipsi- and contralateral projections for all areas studied (Fig. 14E). This poor correlation of ipsi- and contralateral targeting presents a somewhat different picture of circuit function to the corticocortical and corticostriatal inputs (Fig. 3 and 7).

**Figure 14.**
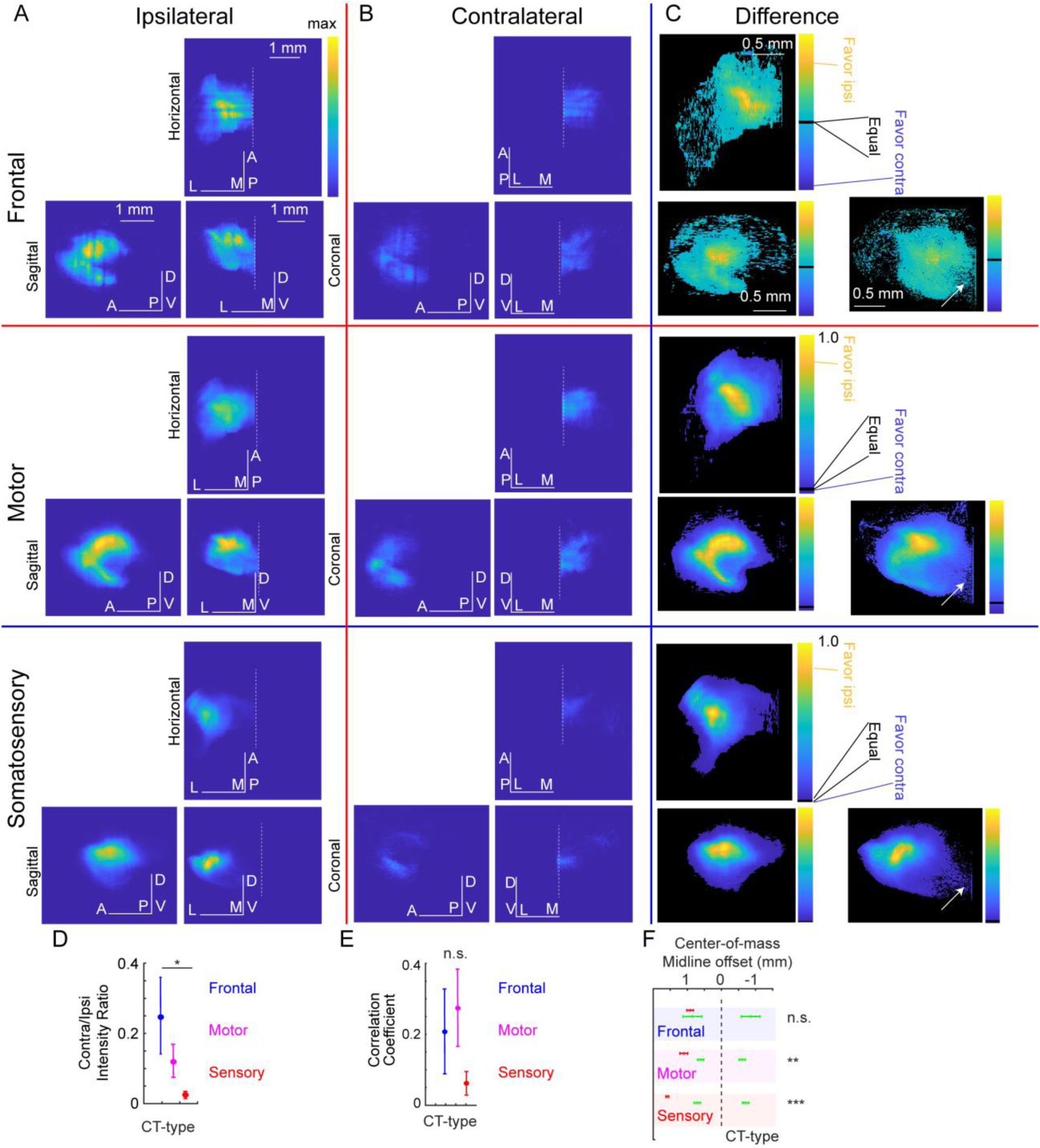
Comparison of ipsi- and contralateral CT-type corticothalamic projections across frontal, motor, and somatosensory cortex. (A) A 3D map of CT-type projections to the thalamus was generated from each injection by selecting only thalamic voxels and nomalized to the peak. These were then averaged with similar injection sites (frontal injections in the top row, motor injections in the second row, and somatosensory injections in the last row). The averaged normalized voxels for the left (ipsilateral) injection site were then displayed as projections along the horizontal (top), sagittal (left) or coronal plane (right). Anterior/posterior (A/P), dorsal/ventral (D/V), and medial/lateral (M/L) as indicated. Midline is shown as a dotted white line in horizontal and coronal planes. (B) The same presentation for the contralateral projections. (C) The difference between ipsilateral and contralateral was computed by subtraction for each brain, then averaged. The subtraction is shown as the left (ipsilateral) view, with black indicating values closest to zero. Arrows in the motor and sensory rows show comparable input where midlnie crossing axons are present. (D) Ratio of intensity of contralateral and ipsilateral projections shown as the mean +/- SE for frontal (blue), motor (purple), and somatosensory (red). (E) Correlation coefficient for ipsi- and contralateral thalamus (note y-axis compared to scores for striatum). (F) Center of mass of ipsilateral (red, left) and contralateral (green, right) thalamic voxels plotted as mean +/- SE from the midline (black dashed line). Contralalateral position plotted in a mirrored location at left for statistical comparison to ipsilateral midline offset. t-test used for all statistical comparions. *p<0.05; **p<0.01; ****; p<0.001.

## Discussion

### Differences in function and somatotopy of cortical areas

Sampling frontal, motor, and sensory areas in an aligned atlas-space provided a cell type-specific dataset for fair comparisons of connectivity using three major areas of neocortex. Somatotopy in S1 is quite precise, with thalamic afferents from VPM targeting single whisker barrels. While contralateral somatosensory information to cortex is limited, callosal corticocortical afferents are present, targeting local cortical circuits in a similar laminar pattern both ipsilaterally and contralaterally (Petreanu et al., 2007). Consistent with this, relatively weak corticocortical and corticostriatal projections exist (Fig. 3 and 7). These afferents are capable of plasticity during sensory manipulation, such as by whisker elimination (Petrus et al., 2020; Petrus et al., 2019), supporting the hypothesis that weak connections may expand in cases where plasticity is adaptive, such as following injury. M1 somatotopy is less precise, with microstimulation mapping imprecisely defining borders (Brecht et al., 2004; Hooks et al., 2011; Li & Waters, 1991; Tennant et al., 2011) and varying considerably between animals (Tennant et al., 2011). This change in precision between S1 and M1 is reflected by differences in striatal somatotopy (Hooks et al., 2018). Remarkably, the precise somatotopy of ipsilateral projections is largely reduced in contralateral striatum (Fig. 7), though some of it is retained in motor areas (Fig 4), perhaps because, while many actions move one limb independently, other movements require symmetrical coordination.

The symmetrical and topographic organization of frontal projections was unexpected, as this area is not previously mapped with the same precision as sensory and motor areas. That frontal cortices may be topographically organized is suggested by the corticocortical connectivity with other somatotopically defined regions of M1 and S1 (Zingg et al., 2014). Microstimulation studies suggest a rostral forelimb area (Tennant et al., 2011), but data for a complete body representation is incomplete. Our data show that frontal IT-type projections to striatal targets are well-organized and reliably project to targets in the ipsi- and contralateral hemispheres.

Projection patterns in ipsilateral cortex (Fig. 3G) and striatum (Fig. 7G) are as sensitive to small displacements in injection location as in somatosensory areas. Furthermore, this topographic precision is maintained in contralateral projections (Fig. 3H and Fig. 7H), making frontal cortex connectivity highly symmetric. There did not seem to be a large difference in contralateral symmetry between medial and lateral sites (ALM versus M2; Fig. 7F), suggesting the gradient of topographic precision in the contralateral projections varies mostly along the anterior-posterior axis.

Primary visual cortex might be similar to S1, with weak callosal connectivity, as the principal function is to represent the contralateral visual field. As incoming acoustic information from a single location might reach both ears, there would be value in interhemispheric connectivity in auditory cortex. Further, some higher order areas in parietal cortex that represent more abstract concepts might be interesting to evaluate similarly.

### Organization and strength of the corticostriatal and corticocortical projection

Our data quantify the substantial contralateral corticostriatal and corticocortical projections of IT-type pyramidal cells. Projections from non-cell type specific injections in wild- type mice (Oh et al., 2014) were qualitatively similar, suggesting our results are not due to assessing a small subset of neurons. The contralateral striatal projection from frontal areas was strong, ∼65% the strength of the ipsilateral projection (Fig. 7D-E). This was considerably weaker for M1 (∼25%) and S1 (∼15%). Cortex had a similar pattern of contralateral cortical projections, though the values were somewhat closer (Frontal, ∼37%; M1, ∼27%; S1, ∼17%). Crossed projections are also stronger from frontal and cingulate areas than from S1 in non-human primates (Borra et al., 2022). Further, the literature suggests that the crossed corticocortical connection instead of the direct corticostriatal one is more effective in causing sensory responses to ipsilateral touch (Reig & Silberberg, 2016). Retrograde tracer injections in striatum label premotor frontal cortex relatively symmetrically while barely labeling contralateral parietal cortex (Borra et al., 2022). Primate posterior parietal cortex (Area 7) also projects contralaterally to topographically similar striatal targets as ipsilateral afferents, though less extensively (Cavada & Goldman-Rakic, 1991). Along the rostro-caudal gradient of the striatum, the caudate head is more likely to receive converging input from ipsi- and contralateral cortex, while the tail is more exclusively ipsilateral (Griggs et al., 2017). Although a simplified interpretation of the data suggests a rostral-caudal gradient in contralateral connectivity, midline crossing projections from associative cortices (frontal and posterior parietal) suggests a functional gradient: regions involved in sensory processing are less likely to have midline crossing projections, while cortical areas related to abstract functions (categorization and sensory-motor coordination) have stronger contralateral connectivity to integrate information and act as a single unit. However, data from rat sensory areas (Alloway et al., 2006) suggests a reasonably strong contralateral cortical projection based on retrograde tracing from striatum.

The main feature of corticostriatal organization is the convergence of excitatory input (including thalamic input) from many afferents onto striatal projection neurons, which in turn project to the indirect pathway (to external globus pallidus) or the direct pathway (to substantia nigra pars reticulata and internal globus pallidus), with some convergence at each step, eventually outputting to thalamus and brainstem (Lee et al., 2020). Our data tests the concept that cortical output generally forms largely parallel, functionally segregated loops from cortex through basal ganglia (Alexander et al., 1986) by including contralateral projections.

### Convergence of ipsilateral and contralateral projections in the striatum

These data reveal a relatively strong and remarkably symmetric organization of the premotor frontal output to striatum. Both medial and lateral frontal areas (M2 and ALM) show strong correlation in their projections to ipsilateral and contralateral striatum. Small injection site displacements shift the targeting of projections (Fig. 7G-I). That these correlations are as precise as S1 suggests that topographic representation in frontal areas exists, though whether this is motor-related or decision-related is unclear. Whether cell type-specific targeting differs for ipsi- and contralateral targets (Johansson & Silberberg, 2020) was also tested for M1 using physiological means, but differences for direct and indirect pathway connectivity were not pronounced. Similar tests have not been performed for frontal projections.

For executive function in premotor frontal areas, connectivity may permit input between hemispheres, helping converge on a single decision for the animal. The motor plan can be stored in one side of frontal cortex during unilateral silencing (and restored to the other side following relief), suggesting that these indeed do operate in concert (Guo et al., 2014; Li et al., 2016). Thus, the purpose of well-organized, near symmetrical projections from frontal areas may be to instantiate a unitary sense of motor planning and executive function, and to use the striatal projections to perform the same computation on incoming information in parallel, potentially supporting motor planning that is robust to noise and perturbation in the network.

### Absence of symmetry in the corticothalamic projection

Corticothalamic projections include two cell types which differ in anatomy and function (Sherman & Guillery, 2006; Veinante et al., 2000). Our study shows that both target contralateral thalamus, with contralateral PT-type projections weaker than CT-type ones. In contrast to corticostriatal and corticocortical axons, corticothalamic inputs from both cell types lacked substantial symmetry (Fig. 13E and 14E). As the thalamocortical output is exclusively ipsilateral (Sherman & Guillery, 2006), integration of premotor frontal cortex computations across the two hemispheres must mainly occur through corticocortical and corticostriatal circuits.

### Comparing the role of rodent and primate premotor areas

In rodents, inhibition of ALM biased movement to the ipsilateral side. However, IT-type neuron activity in ALM has been shown to split roughly equally between ipsi- and contra- movements, with most cells responding jointly to movement and preparatory phases of the task (Li et al., 2015). This suggests IT-type cells provide a planning/preparatory signal. Our findings of bilateral symmetry in corticocortical and corticostriatal projections from ALM IT-type cells support this hypothesis and help to explain the lack of contralateral selectivity in ALM IT-type cells. A similar pattern of bilaterally symmetric connectivity to dorsal striatum and motor cortex is present in human dorsal premotor cortex (Hardwick et al., 2015; Ruddy, Leemans, & Carson, 2017). Transcallosal corticocortical connectivity varied between cortical areas, with dorsal premotor cortex and supplementary motor areas having the densest contralateral connectivity (Ruddy, Leemans, & Carson, 2017), suggesting these areas bridge executive cognitive functions and motor control. It is unclear which primate premotor areas (Dum & Strick, 2002) are the closest homologues to rodent premotor ones. The dorsal premotor cortex (Area 6) shares some similarity in connectivity and function. It is reciprocally connected to M1 and is tuned to sensory-guided movements (Geyer et al., 2000). Further, the rodent premotor areas studied here (ALM and M2) contain PT-type neurons, akin to the pyramidal tract contributions of dorsal premotor and supplementary motor areas.

Consistent with our rodent results, primate corticostriatal projections are bilateral and have higher density on the ipsilateral side. Contralateral striatal projections are extensive for premotor areas and weaker, but present, in M1 (Kunzle, 1975, 1978). Similar to our weaker S1 contralateral projections, primate S1 does not appreciably project to contralateral striatum (Flaherty & Graybiel, 1991; Jones et al., 1977).

## Conclusions

What is the role of such contralateral connectivity? Under normal conditions, callosal projections may mediate interhemispheric inhibition, where projections suppress responses in the opposite hemisphere, isolating processing that should be unilateral (Carson, 2020; Chiarello & Maxfield, 1996), though the relative effect of excitation and inhibition is better tested with cell- type specificity in rodent models of callosal connectivity (Anastasiades et al., 2018).

Functionally, contralateral connectivity supports stroke recovery (Hensel et al., 2023) and improves motor learning on the side contralateral to the trained limb (Ruddy, Leemans, et al., 2017). Here, we suggest that, in addition to plasticity, cortical areas that engage in motor planning – where it is crucial that the brain converge on a single motor plan – must share the output of their computation via crossed pathways, which likely include the crossed corticocortical and corticostriatal pathways described here. Maintaining some degree of symmetry helps achieve integration of circuit components representing the same aspect of motor planning in these regions.

## Acknowledgements

We thank Afonso Silva, Rob Turner, Andreea Bostan, and members of the Hooks lab for comments and suggestions. We thank Jack Glaser, Paul Angstman, Sue Tappan, Mike Fay and Scott Gerfen at MBF Bioscience for the collaboration to develop and use the BrainMaker software. This work was supported by a NARSAD Young Investigator Award (BMH), NIH NINDS R01 NS103993 (BMH), a CDMRP PRMRP Discovery Award PR201842 (BMH), a Whitehall Foundation award (BMH), and a National Institute of Mental Health IRP award (ZIA MH002497-33) to C.R.G.

## Author contributions

A.E.P, B.M.H., and C.R.G. conceived of the project and analyzed the data. A.E.P, M.H., N.J.O., and B.S.E. developed software for analyzing the data. R.F.P. and C.R.G. sectioned and imaged the tissue. B.M.H., A.E.P., R.S.W., R. P. S., and C.R.G wrote the paper with help from all authors.

